# Identification of the Regulatory Elements and Protein Substrates of Lysine Acetoacetylation

**DOI:** 10.1101/2024.10.31.621296

**Authors:** Qianyun Fu, Terry Nguyen, Bhoj Kumar, Parastoo Azadi, Y. George Zheng

**Affiliations:** Department of Pharmaceutical and Biomedical Sciences, College of Pharmacy, University of Georgia, Athens, Georgia, USA; Complex Carbohydrate Research Center, University of Georgia, Athens, Georgia, USA

**Keywords:** acyltransferase, epigenetics, histones, ketone body, lysine acylation, metabolism

## Abstract

Short chain fatty acylations establish connections between cell metabolism and regulatory pathways. Lysine acetoacetylation (Kacac) was recently identified as a new histone mark. However, regulatory elements, substrate proteins, and epigenetic functions of Kacac are not yet fully understood, hindering further in-depth understanding of acetoacetate modulated (patho)physiological processes. Here, we created a chemo-immunological approach for reliable detection of Kacac, and demonstrated that acetoacetate serves as the primary precursor for histone Kacac. We report the enzymatic addition of the Kacac mark by the acyltransferases GCN5, p300, and PCAF, and its removal by deacetylase HDAC3. Furthermore, we establish acetoacetyl-CoA synthetase (AACS) as a key regulator of cellular Kacac levels. A comprehensive proteomic analysis has identified 139 Kacac sites on 85 human proteins. Bioinformatics analysis of Kacac substrates and RNA-seq data reveal the broad impacts of Kacac on multifaceted cellular processes. These findings unveil pivotal regulatory mechanisms for the acetoacetate-mediated Kacac pathway, opening a new avenue for further investigation into ketone body functions in various pathophysiological states.

## Introduction

Cellular metabolites play crucial roles in the production of bioenergy and the synthesis of biomolecules (1). In addition to this classical functionality, there is a growing acknowledgment that certain metabolic molecules also exhibit regulatory functions by acting as precursors for post-translational modifications (PTMs) of proteins (2). Short-chain fatty acids are abundant metabolites widely present in both prokaryotic and eukaryotic cells (3). Fatty acylation of lysine residues in proteins, acetylation being the most prominent, has been known as a versatile form of reversible PTMs for their capacity to impart diverse regulatory functions on key cellular processes (4). Lysine acylation reactions are dependent on their respective short-chain CoAs as cofactors (more precisely termed as cosubstrates) and are regulated by a finely balanced enzymatic counteraction involving lysine acetyltransferases (KATs) and lysine deacetylases (KDACs) (5,6). Beyond lysine acetylation, in recent years, multiple fatty acylations on nuclear histones, e.g. lysine β-hydroxybutyrylation (Kbhb) (7), isobutyrylation (Kibu) (8), have been identified, indicating complex connections between cellular metabolism and epigenetic regulation of gene expression. Of note, while most fatty acylations were initially identified on nuclear histones, it is later found that reversible acylation serves as a significant regulatory mechanism for a diverse range of cellular proteins across multiple cellular compartments (9). Indeed, growing evidence suggests a strong correlation between short-chain lysine acylations and diverse pathophysiological conditions, contributing to the progression of diseases (10–12).

Ketone bodies, including acetoacetate (AcAc), D-β-hydroxybutyrate (BHB) and acetone, are produced in the liver and subsequently distributed throughout the body during extended periods of dietary carbohydrate restriction and fasting. Apart from serving as an energy source, ketone bodies recently are starting to be acknowledged for their roles as signaling mediators, modulators of inflammation and oxidative stress (13). Their importance extends to critical pathological conditions that span cardiovascular, cerebrovascular, neurological disorders such as failing hearts (14), ischemic stroke (15), Parkinson’s disease, Alzheimer’s disease (16,17) and cancers (18,19). The Zhao group first found that β-hydroxybutyrate acts as a precursor for covalent modification of histones in the form of lysine β-hydroxybutyrylation (Kbhb) (7). In the mice liver subjected to fasting or streptozotocin-induced diabetes, Kbhb sets up a connection between chromatin regulation and the cellular pathophysiological functions of β-hydroxybutyrate (7). Further studies investigated key regulatory elements and substrate specificity of Kbhb (20). The β-hydroxybutyrate-mediated Kbhb of p53 pinpoints the connection between Kbhb and tumorigenesis (21). A recent study highlights that acetoacetate, another ketone molecule, also acts as a precursor for an uncharacterized histone PTM mark, known as lysine acetoacetylation (Kacac) (22). Nevertheless, key regulatory elements controlling Kacac formation, downstream targets of this modification at the proteomic level, as well as possible functional impacts of lysine acetoacetylation on cellular processes, remain as mysterious subjects of investigation for the field. These problems are particularly urgent to be addressed in order to gain a clear understanding of the regulatory mechanism of acetoacetate, especially considering its distinct function compared to β-hydroxybutyrate.

As a new PTM biomarker, there lacks effective technical tools to study Kacac in proteins. In recent years, the combination of specific antibody-based immunoprecipitation with advanced high-resolution mass spectrometry (MS) has streamlined the identification of PTM marks on proteins (6). Nevertheless, the methods for generating new antibodies targeting small antigen markers such as protein acylation are typically laborious, time-intensive, expensive, and of low success rate. In this study, we introduce a distinctive and dependable chemo-immunological approach, wherein Kacac is first reduced into the Kbhb marker using NaBH_4_ and subsequently detected using a currently commercially available anti-Kbhb antibody. This innovative method circumvents the necessity to develop new antibodies for Kacac, and meanwhile allows for simultaneous measurement and comparison of Kacac with Kbhb marks on proteins. Through implementing our developed chemo-immunological method, we present the systematic profiling of the Kacac mark as a novel, acetoacetate-induced modification in both histone and non-histone proteins in the human proteome. Our proteomic screen identified 139 unique Kacac sites across 85 proteins in HEK293T cells. Detailed bioinformatics analysis and RNA-seq results reveal that Kacac has unique physiological significance and potentially participates in a balanced system with Kbhb to co-regulate cellular pathways. Expanding upon previously reported writers and erasers, we have identified p300, GCN5, and PCAF as acetoacetyltransferases in vitro, and HDAC3 as a de-acetoacetylase in vivo. Furthermore, we illustrated the mechanism behind acetoacetate-induced histone Kacac through enzymatic conversion by acetoacetyl-CoA synthetase (AACS). Together, this study delves into the complex mechanisms of ketone body-mediated Kacac, broadening the list of protein substrates and pathways potentially regulated by Kacac. The findings provide a foundational understanding for further exploration of the dynamic control mechanisms of ketone body homeostasis and the potential downstream roles of ketone bodies in regulating human disease processes.

## Results

### Method development for the detection of lysine acetoacetylation

Lysine acetoacetylation is a novel PTM and currently there is no commercial antibody to study Kacac. To streamline the process of specific detection of Kacac for either immunoblotting or immunoprecipitation applications, we attempted to develop a reliable chemo-immunological approach for the detection of Kacac. Specifically, following introduction of lysine acetoacetylation on proteins, the 3-carbonyl group in the Kacac marker was converted to a hydroxyl group using the reducing reagent NaBH_4_ under slightly basic condition. This conversion transforms the Kacac mark into β-hydroxybutyryllysine (Kbhb), which can then be detected using commercially available and widely used anti-Kbhb antibody. A big advantage of this chemo-immunological approach is that it enables the simultaneous detection of Kbhb and Kacac under various experimental conditions, and further allows for the identification and distinction of the Kacac mark from Kbhb through proteomic analysis if deuterated precursors are used (Fig. 1A). To assess the practicability of our proposed method, we synthesized a peptide containing H2B N-terminal sequence with an acetoacetyl group at Lys-15 position, i.e., H2B(1–26)K15acac (Supplementary Fig. S1). This is a Kacac site identified from our preliminary data. The acetoacetylated peptide was reduced using NaBH_4_ under basic conditions, with a parallel sample lacking NaBH_4_ under the same conditions serving as the control. The specificity of the pan anti-Kbhb antibody was then assessed via a dot blot assay (Fig. 1B). In this assay, the pan-anti-Kbhb antibody demonstrated an excellent performance for the recognition of the Kacac peptide only after NaBH_4_ reduction, supporting the feasibility of our developed chemo-immunological method. We want to point out that the pan-Kbhb antibody used herein is independent of peptide sequence so different modification sites on proteins shall all be recognized. Therefore, we subsequently employed this technical strategy to delve into the biology of lysine acetoacetylation.

**Fig. 1.**
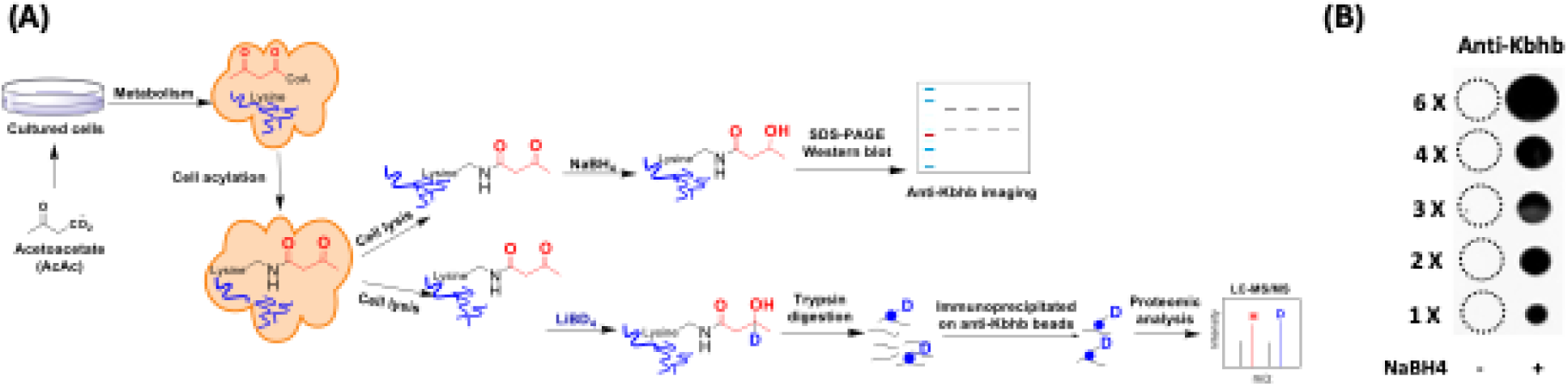
Methods development for the identification of lysine acetoacetylation. **(A)** Biosynthetic pathways for lysine acetoacetylation (Kacac) and proposed detection methods of Kacac. **(B)** Dot blot assay verifying the specificity of the pan anti-Kbhb antibody in our developed methods. Synthetic H2BK15acac peptide (Ac-PEPAKSAPAPKKGSKacacKAVTKAQKKDG-NH2) was used for the assay.

### Acetoacetate is a precursor source for histone Kacac generation in cells

Recent studies have unveiled that short-chain fatty acids prompt the production of their respective acyl-CoAs within cells, consequently enhancing levels of histone acylations (23). We posit that acetoacetate enters the cell and is converted into acetoacetyl-CoA to serve as the cofactor for generating Kacac marks in cellular proteins. The recent study by Zhao and coworkers incubated HepG2 cells with ethyl acetoacetate and reported a dose-dependent increase in histone Kacac upon treatment, as detected by a specifically designed antibody (not commercially available) (22). To directly prove that acetoacetate induces Kacac levels, we cultured HEK293T cells with the treatment of lithium acetoacetate at different concentrations (0, 5, 10, 20 mM). Following treatment, we extracted the nuclear histone proteins, which were then reduced using NaBH_4_. The proteins were separated on a 15% SDS-PAGE gel, subsequently transferred to a nitrocellulose membrane, and probed using an anti-Kbhb antibody. Western blotting of histones revealed faint band intensities in the absence of NaBH_4_ treatment, indicating the presence of endogenous Kbhb levels on the histones (Fig. 2A). However, after reduction with NaBH_4_, the intensity of the western blot bands increased substantially in an acetoacetate dose-dependent manner on the core histones. Clearly, this augmentation in western blot intensity was attributed to the heightened levels of Kacac resulting from acetoacetate treatment. Similarly, we observed an elevation in global histone Kacac levels in HCT116 cells, further supporting that acetoacetate served as the precursor for acetoacetylation (Fig. 2B). We also examined the time course of acetoacetate-induced Kacac and found that histone Kacac became detectable as early as 1 h following acetoacetate treatment (Supplementary Fig. S2A). Notably, this chemo-immunological method has the advantage of allowing for the concurrent assessment of endogenous Kbhb levels and acetoacetate-induced Kacac levels on histones (Figs. 2A, 2B). Our results indicated a subtle increase in histone Kbhb following acetoacetate treatment, suggesting that acetoacetate may play a minor role in promoting Kbhb generation within the cell.

**Fig. 2.**
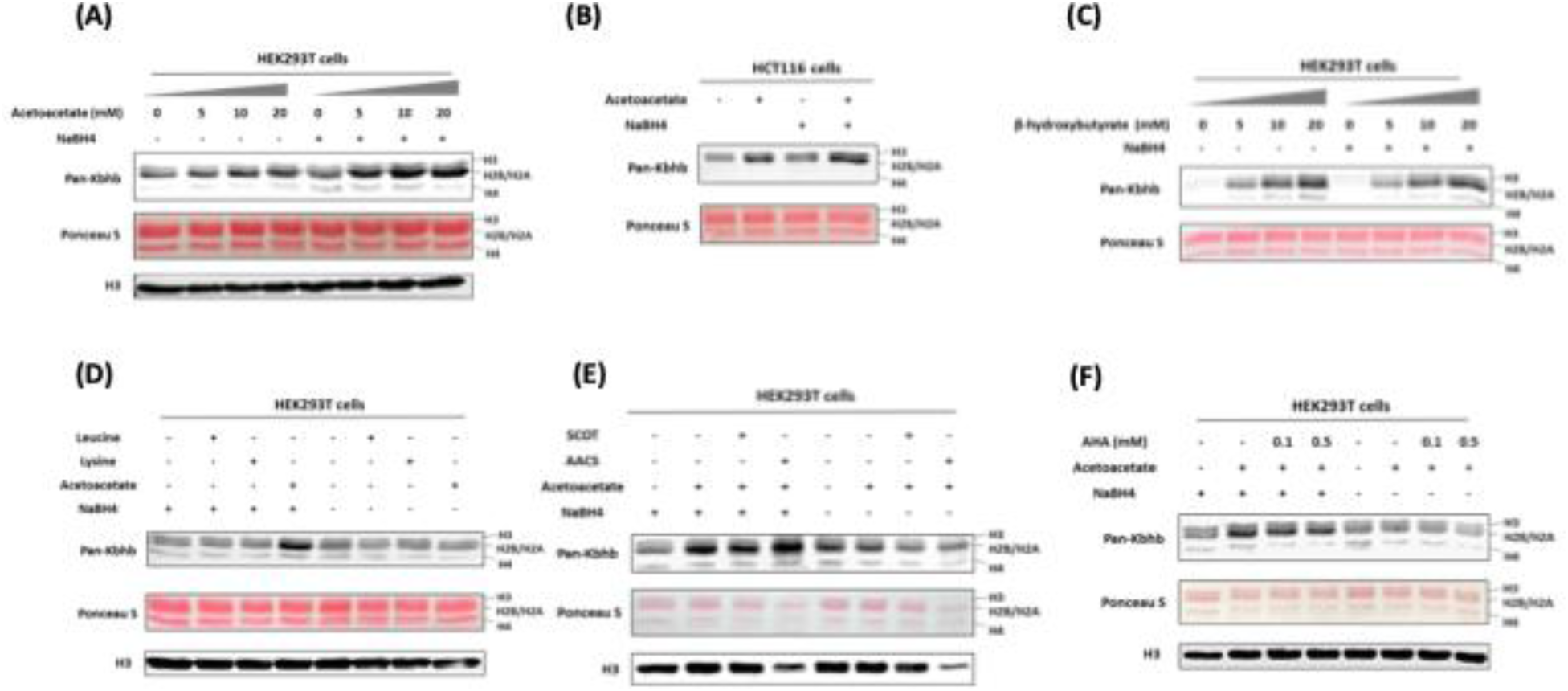
Acetoacetate dynamically regulates Kacac levels through the generation of acetoacetyl-CoA. **(A)** Western blot analysis of histones from HEK293T cells treated with increasing doses of lithium acetoacetate. **(B)** Detection of histone Kacac in HCT116 cells. **(C)** Western blot analysis of histones from HEK293T cells treated with increasing doses of sodium β-hydroxybutyrate. **(D)** Western blot analysis of histone Kacac in response to treatment with lithium acetoacetate or ketogenic amino acids (leucine and lysine) in HEK293T cells. **(E)** Western blot analysis of histone Kacac in response to overexpression of ketolysis enzymes (SCOT and AACS) in HEK293T cells. **(F)** Western blot analysis of histone Kacac in response to treatment with acetohydroxamic acid (AHA), a known SCOT inhibitor, in HEK293T cells.

To gain more insights into the dynamics of histone Kacac stimulated by acetoacetate, we treated HEK293T cells with other metabolic molecules that may influence Kacac levels within cells. We first investigated whether β-hydroxybutyrate could elevate histone Kacac levels in cells. By comparing the band intensities before and after NaBH_4_ treatment, no increase in Kacac levels was observed following β-hydroxybutyrate treatment of HEK293T cells (Fig. 2C). This suggests that β-hydroxybutyrate did not affect Kacac levels in HEK293T cells. To delve deeper into the precursor sources for histone Kacac generation, we examined the impact of two ketogenic amino acids—leucine and lysine—on Kacac levels. As shown in Fig. 2D, only acetoacetate treatment caused Kacac increase, neither leucine nor lysine had an ability to increase Kacac levels on histones. These results indicated that acetoacetate acts as a predominant metabolic precursor for histone Kacac in the tested cells.

### AACS enzyme dynamically regulates histone Kacac in cells

It is known that acetoacetyl-CoA synthetase (AACS) specifically catalyzes the activation of acetoacetate to its coenzyme A ester and serves as a highly regulated lipogenic enzyme in numerous tissues involved in *de novo* lipid synthesis (24). We conjecture that exogenous acetoacetate elevates histone Kacac levels through directly generating acetoacetyl-CoA by the AACS. Thus, we investigated whether overexpression of the AACS would enhance histone Kacac levels in cells. We transiently transfected HEK293T cells with flag-tagged AACS plasmid followed by acetoacetate treatment. Immunoblot analysis of total cell lysates proved that AACS was successfully overexpressed at the protein level (Supplementary Fig. S2B). As anticipated, a substantial increase in histone Kacac levels was observed in HEK293T cells upon simultaneous overexpression of AACS and acetoacetate treatment (Fig. 2E), supporting that AACS utilized acetoacetate to generate acetoacetyl-CoA priming for histone acetoacetylation. Succinyl CoA-oxoacid transferase (SCOT) is a mitochondrial enzyme thought to catalyze the regeneration of acetoacetyl-CoA from acetoacetate and facilitate ketone body oxidation for energy production (25). To further assess the potential contribution of SCOT to the acetoacetyl-CoA pool for histone Kacac, we conducted transient transfection to overexpress SCOT in HEK293T cells (Supplementary Fig. S2B). Nevertheless, our results did not reveal any significant increase in histone Kacac levels following SCOT overexpression (Fig 2E). We subsequently examined the effect of a previously described SCOT inhibitor, acetohydroxamic acid (AHA) (26), to validate the impact of SCOT on histone Kacac levels in HEK293T cells. Upon treating cells with AHA, we observed a slight decrease in signals, indicating a subtle effect of SCOT on histone Kacac (Fig. 2F). Drawing from these findings, we conclude that AACS is the primary enzyme contributing to the acetoacetyl-CoA pool for histone Kacac generation in HEK293T cells, whereas SCOT exhibits minimal effects.

In a prior study, it was reported that AACS mRNA exhibited high expression levels in the kidney, heart, and brain, while demonstrating lower expression in the liver (27). However, AACS was strongly upregulated in liver cancer cells, influencing cancer development and progression (28). The varied expression and distinct functions of AACS promoted us to investigate whether it exerts similar effects on histone Kacac levels in HepG2 liver cells. To accomplish this, we transiently overexpressed AACS in HepG2 cells (Supplementary Fig. S2C) and evaluated its impact on Kacac levels using our established methods. Surprisingly, the overexpression of AACS in HepG2 cells did not result in a significant increase in histone Kacac levels following acetoacetate treatment (Supplementary Fig. S2D). We also tested if HMG-CoA reductase (HMGCR) affects Kacac formation. We either overexpressed HMGCR or inhibited its activity using the lovastatin inhibitor in HepG2 cells. Subsequently, we treated the cells with acetoacetate to observe the impact of HMGCR on histone Kacac level. The chemo-immunological western blot results showed HMGCR modulation has little effect on the Kacac level (Supplementary Figs. S2D, S2E).

### p300, PCAF and GCN5 mediate histone acetoacetylation

Mounting evidence suggests that classical lysine acetyltransferases can govern a range of newly discovered acyl modifications (6). To comprehensively identify enzymes involved in mediating Kacac on histones, we sought to express and purify the major human histone acetyltransferases (HATs), including GCN5, PCAF, p300, Tip60, MOF, MOZ, HAT1, MORF, and HBO1. The SDS-PAGE analysis validated both the identity and the purity of these enzymes (Supplementary Fig. S3A). With these recombinant proteins, we tested their activity to acetylate or acetoacetylate recombinant histone proteins through western blot assay. Individual HAT enzymes were incubated with histone H3/H4 and acetoacetyl-CoA in the reaction buffer. The reaction product was detected using anti-Kbhb antibody, with and without NaBH_4_ reduction. While most of these enzymes displayed well-characterized acetyltransferase activities, only p300, PCAF, and GCN5 demonstrated strong histone acetoacetyltransferase activities (Figs. 3A, 3B, Supplementary Figs. S3B-S3E). Interestingly, GCN5 and PCAF exhibited a markedly higher acetoacetyltransferase-to-acetyltransferase ratio compared to that of p300. HAT1 and some MYST members exhibited weak acetoacetyltransferase activity (Fig. 3A, Supplementary Figs. S3C, S3D, S3F). Subsequently, we conducted additional tests to further ascertain whether the acetoacetyltransferase activities of p300, GCN5, and PCAF could be detected on cellular histone substrates. This involved incubating acetoacetyl-CoA and histone extracts from HEK293T cells with p300, GCN5, or PCAF. As expected, stronger Kacac signals were observed in the groups with added enzymes (Supplementary Fig. S3F). To investigate whether p300 regulates histone Kacac in cells, we transiently transfected HEK293T cells with p300 plasmid, followed by treating the cells with acetoacetate. In accordance with the aforementioned observations (Fig. 2), acetoacetate markedly elevated histone Kacac levels (Supplementary Fig. S3G). Unfortunately, the coexistence of p300 overexpression with acetoacetate treatment did not appreciably enhance Kacac levels (Supplementary Fig. S3G). It could be possible that native p300 protein levels in HEK293T cells are quite high which shields the effect of the exogenously expressed proteins.

**Fig. 3.**
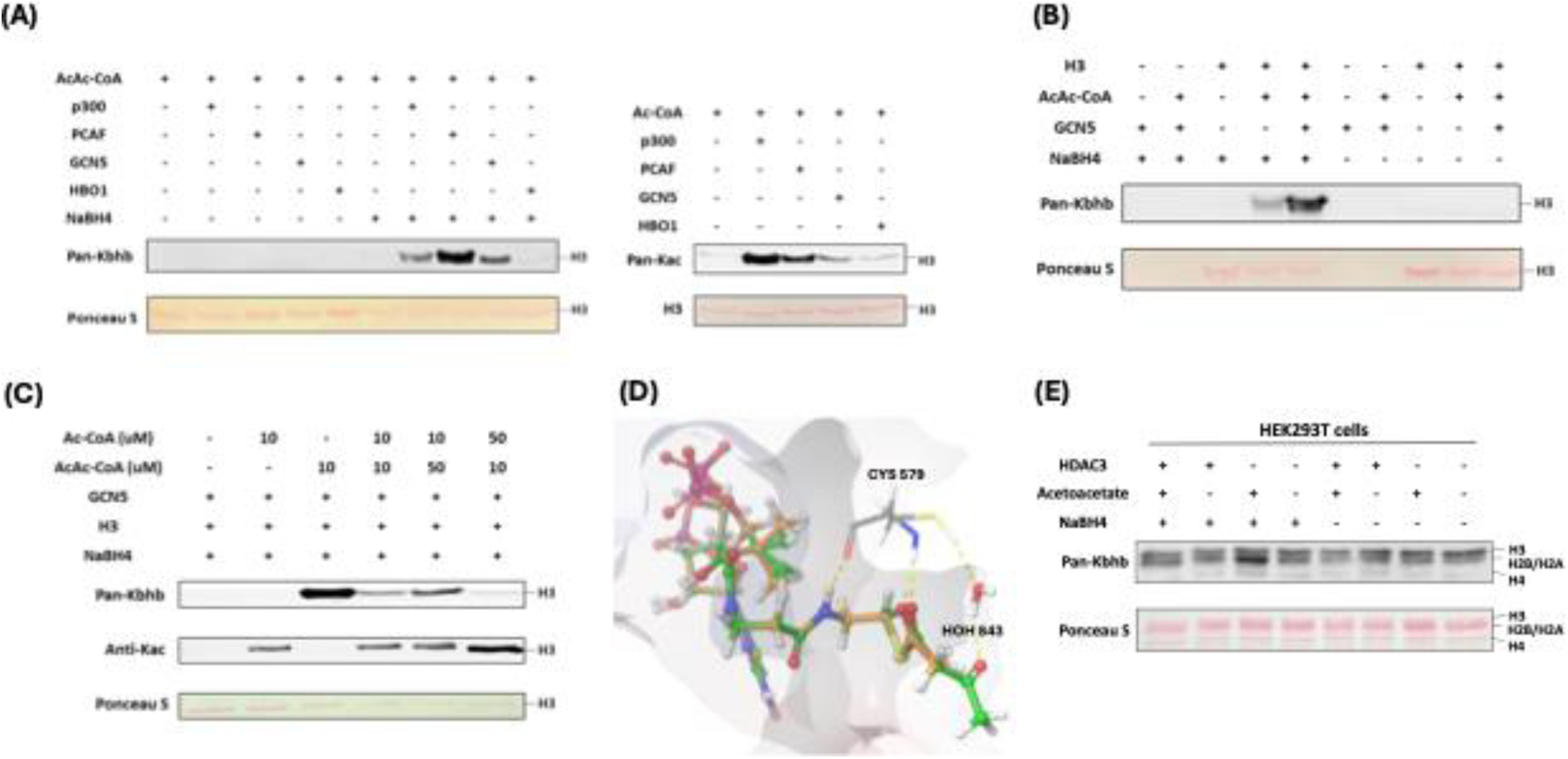
Identification of writers and eraser responsible for regulating histone Kacac. **(A)** p300, PCAF and GCN5 exhibited remarkable acetoacetyltransferase activities (left) and acetyltransferase activities (right) on recombinant histone H3 proteins. **(B)** Validation of GCN5-mediated Kacac on recombinant histone H3 proteins. **(C)** Proportional changes of acyl-CoAs result in dynamics of substrates on recombinant histone H3 proteins. **(D)** Diagram illustrating the catalytic pocket of GCN5 bound with acetyl-CoA (orange) and acetoacetyl-CoA (green). PDB: 5TRL was used for the modeling. **(E)** Overexpression of HDAC3 abolished acetoacetate-induced Kacac in HEK293T cells.

Next, we incubated recombinant p300, PCAF, or GCN5 with acetoacetyl-CoA and synthetic histone peptides under identical HAT assay conditions. The resulting reaction mixture was subjected to MALDI-MS analysis, which revealed expected product peaks (M+84) in all the catalytic reactions, thus providing further confirmation of the acetoacetyltransferase activities of the tested enzymes (Supplementary Fig. S4A).

Given that p300, GCN5 and PCAF can catalyze both histone Kac and Kacac, we predicted that alterations in a modified substrate would be directly influenced by fluctuations in the concentrations of acetyl-CoA or acetoacetyl-CoA within cells. These, in turn, would be closely linked to the regulation of these cofactors originating from both inside and outside the cell (29). To gain insights into how acylations are differentially regulated in mammalian cells, we performed competition experiments by incubating recombinant histone H3 substrate with the HAT enzymes we tested above and varying ratios of acyl-CoAs. Indeed, altering the relative concentrations of acetyl-CoA or acetoacetyl-CoA showed differential impacts on the proportion of each modification present in the final reaction product. Acetyl-CoA was a superior cosubstrate of p300, GCN5, and PCAF. Presence of acetoacetyl-CoA did not affect acetyl-CoA dependent histone acetylation (Fig. 3C, Supplementary Figs. S4B, S4C). However, H3 acetoacetylation was significantly suppressed by the presence of acetyl-CoA, whereas increasing concentrations of acetoacetyl-CoA led to elevated Kacac levels. p300 was previously reported to catalyze the enzymatic addition of β-hydroxybutyryl group to lysine (20). To directly compare p300’s activity as a β-hydroxybutyryltransferase or acetoacetyltransferase, we conducted additional incubations using recombinant human histone proteins along with varying ratios of acetoacetyl-CoA and β-hydroxybutyryl-CoA. Surprisingly, the presence of acetoacetyl-CoA scarcely influenced the generation of β-hydroxybutyrylation, suggesting that p300 exhibits stronger activity as a β-hydroxybutyryltransferase than an acetoacetyltransferase (Supplementary Fig. S4D).

To further understand the acetoacetylation activity of HATs described above, we conducted a silico molecular modeling of acetoacetyl-CoA/acetyl-CoA with HATs to explore potential binding interactions between acetoacetyl-CoA and the individual HATs. In the structure of GCN5 (PDB ID: 5TRL), the co-crystallized ligand is succinyl-CoA (30) and the carbonyl group of the succinate proved useful in validating the positioning of the acyl groups in the docked acetoacetyl-CoA and acetyl-CoA binding. Several residues were found to interact with both succinyl-CoA and the docked acetoacetyl-CoA and acetyl-CoA: Cys579, Val581, Val587, Gly589, Gly591, Thr592, Lys624, and Tyr617. The hydrogen bonding between the backbone of Ala614 and the co-crystallized succinyl-CoA is lost upon docking of acetyl-CoA and acetoacetyl-CoA. However, the catalytic water in the binding pocket retains its interactions with both acyl-CoAs. We observed that the distance between the sulfur of the co-crystallized succinyl-CoA and the thiol of Cys579 was maintained in the docking poses for the two acyl-CoAs with a distance of 6.3 Å for all three CoA derivatives (Fig. 3D, Supplementary Fig. S5A). In PCAF (PDB ID: 4NSQ), the co-crystallized ligand was CoA (31), many of the interactions found with this molecule were shared with the acyl-CoAs such as Cys574, Val576, Val582, Gly584, Thr587, Lys619, and Tyr612. Acetoacetyl-CoA was found to have more steric clashes towards the terminal acetyl group whereas acetyl-CoA seems to have a favorable hydrogen bonding interaction with Asp610. Here, the sulfurs of the CoA derivatives were found at the following distances from the catalytic Cys574: 3.43 Å for CoA, 3.59 Å for acetyl-CoA, and 3.34 Å for acetoacetyl-CoA. The western blot data suggests that acetyl-CoA may be a better substrate for PCAF (Supplementary Fig. S4B). This might be explained by the orientation of the acetyl group displayed in the modeling when compared to the acetoacetyl group. The carbonyl in the acetyl-CoA is in a position more similar to the Bürgi-Dunitz angle with respect to thiol of Cys574 making the carbonyl more available for nucleophilic attack whereas acetoacetyl-CoA has the carbonyl reversed with its oxygen pointing towards the thiol (Supplementary Fig. S5B). For p300 (PDB ID: 5LKU), compared to the co-crystallized CoA in the structure (32), the interactions with Leu1398, Ser1400, Thr1411, Trp1466, and Ile1457 are maintained when docked with acetyl-CoA and acetoacetyl-CoA. With acetyl-CoA, the pi-cation interaction with Arg1462, observed in the co-crystallized CoA, is maintained. However, the larger acyl group in acetoacetyl-CoA causes a slight shift in the adenyl ring, resulting in the loss of this interaction. Most importantly, the 3’ phosphate in the acyl-CoAs is shown to interact with Arg1410 which is typical for CoA binding (33). The catalytic Cys1438 displays a distance of 6.61 Å between the side chain and the sulfur of CoA as well as acetyl-CoA and 6.64 Å for acetoacetyl-CoA (Supplementary Fig. S5C).

### HDAC3 is a lysine deacetoacetylase in cells

In recent years, certain histone deacetylases (HDACs) have been reported to display non-canonical enzymatic activities towards various protein acylations, including de-succinylation (34), de-β-hydroxybutyrylation (20), and others.

In a recent screening of individual HDACs using synthetic Kacac peptides, it was discovered that HDAC3 demonstrated distinct de-acetoacetylation activity in vitro (22). To investigate whether HDAC3 exhibits bona fide de-acetoacetylation activity in vivo, we conducted HDAC3 overexpression experiments in HEK293T cells through transient transfection. Then, we examined histone Kacac levels using our developed chemo-immunological method. Intriguingly, the overexpression of HDAC3 in HEK293T cells counteracted the increase in Kacac induced by acetoacetate treatment (Fig. 3E). These findings support that HDAC3 functions as a deacetoacetylase both in vitro and in vivo.

To rationalize the de-acetoacetylation activity of HDAC3, small-molecule substrate mimics containing acyllysine (acetyllysine mimic or acetoacetyllysine mimic) were docked into the crystal structure of HDAC3 (Supplementary Fig. S5D) (35). In these molecules, lysine was alpha-N acetylated and C-amidated, so they more closely resemble the backbone of a protein rather than that of a standalone amino acid. The proposed binding mode positions the side chain nitrogen of acetyl lysine within 3.86 Å of zinc and 3.08 Å of Tyr298 while acetoacetyl lysine has its nitrogen within 4.50 Å of zinc and 4.24 Å of Tyr298. Additionally, both docked acyl lysines maintain hydrogen bonding with His134. The poses for both molecules coincide with the methionine model presented by Watson et al, which can serve as a validation for the lysine substrate (35). When comparing the XP GScores, the acetoacetyl lysine mimic docking pose displayed a GScore of -8.899, while the acetyl lysine mimic had a GScore of -7.255. This may be because the docking model for the acetyl lysine mimic shows a steric clash between the methyl group of the acetyl and Cys145. In contrast, the acetoacetyl lysine mimic appears to bind further into the pocket, avoiding this steric interaction. This computational model provides a structural insight into the de-acetoacetylation activity of HDAC3.

### Proteome-wide identification of Kacac substrates in human cells

To understand the scope and impact of lysine acetoacetylation, we mapped out the landscape of Kacac substrates in the HEK293T cell proteome. To do so, the cellular proteome was extracted and reduced using LiBD_4_, followed by trypsin digestion. The resulting tryptic peptides were enriched using anti-Kbhb antibody-conjugated resin. The enriched peptides were then released from the resin for HPLC-MS/MS analysis and database searching. We identified 24 histone Kacac modification sites located at H1, H2A, H2A.Z, H2B, H3 and H4: 10 were the same as the previously reported and 14 were new sites (Fig. 4A). Additionally, our methods enabled the successful identification of 139 sites on 85 cellular proteins (Fig. 5A, Supplementary Table S1). Among these Kacac (DKbhb) proteins, 60 proteins (70%) contain a single Kacac site, 13 proteins (15%) contain two Kacac sites, and the rest contain three or more Kacac sites (Fig. 5A). Impressively, nucleophosmin (NPM) contains 4 Kacac sites, nucleolar and coiled-body phosphoprotein 1 (NOLC1) protein and histone H2B protein bear 5 Kacac site, histone H3 protein bears 6 sites, and nucleolin (NUCL) protein bears 10 sites. It is worth emphasizing that in this experiment, the Kacac marks were identified as the deuterated Kbhb (DKbhb) form. Importantly, endogenously existing Kbhb was also detected from this single experiment. Therefore, our chemoimmunological method, combined with MS/MS, enables the simultaneous identification of both Kacac and Kbhb sites. To examine potential Kacac motifs in the identified substrate proteins, we compared the amino acid sequences surrounding Kacac sites against the background of the human proteome. The data revealed a preference for Ser, Ala or Gly at the -1 and +1 positions. Additionally, proline was highly enriched at the -2 positions while Leu was largely depleted at most positions. Interestingly, a significant preference was observed for alanine and positively charged lysine at multiple positions (-6, -5, -4, 1, 2, 3, 4, 5, and 6), whereas negatively charged amino acids (Asp, Glu) were consistently underrepresented at the -1 and -6 positions (Fig. 5B). These features closely align with the reported flanking sequence preference of Kbhb but differ from the motif analysis results of lysine acetylation (Kac), crotonylation (Kcr), malonylation (Kmal), succinylation (Ksucc), and 2-hydroxyisobutyrylation (Khib) (9,36–40).

**Fig. 4.**
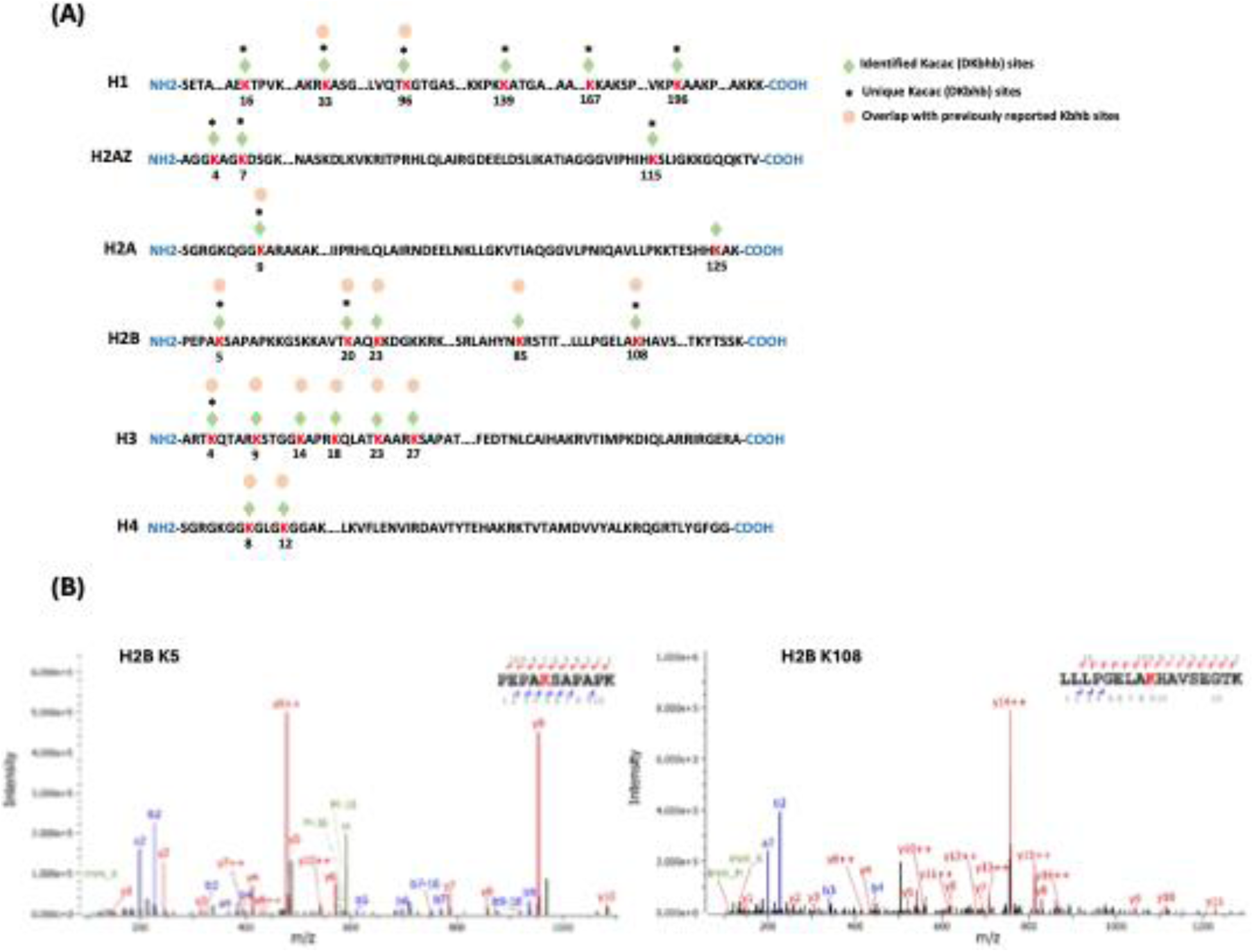
Proteomic screening of histone Kacac sites in HEK293T cells. **(A)** Illustration of histone DKbhb (Kacac) sites identified in HEK293T cells. Green diamond indicates Kacac sites detected in our study. * denotes previously unknown histone Kacac sites. For comparison, the overlapped known Kbhb sites (labeled with an orange dot) described in the literature are also listed. **(B)** MS/MS spectra of two representative DKbhb peptides derived from HEK293T histones.

**Fig. 5.**
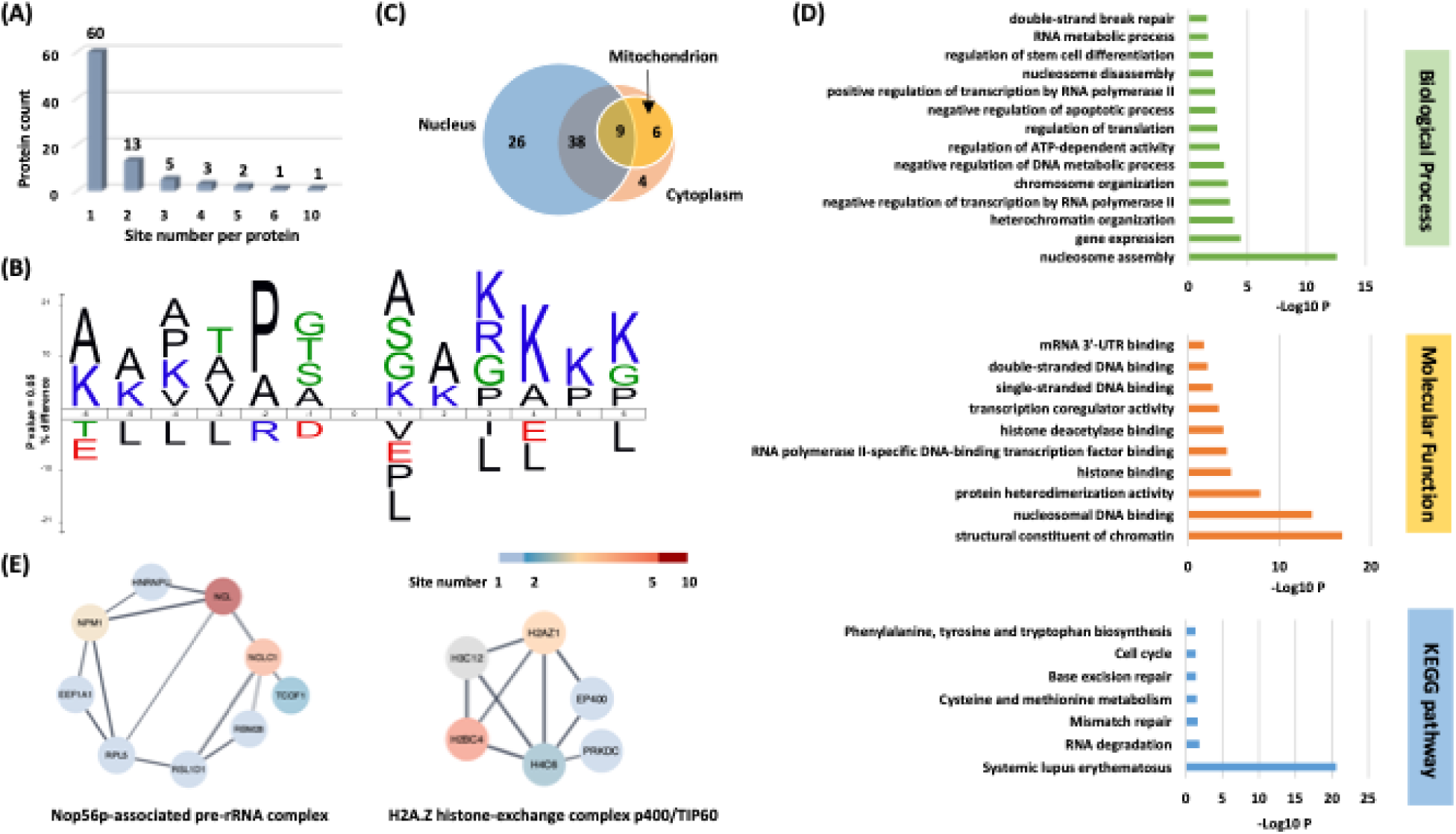
Systematic profiling of the Kacac proteome. **(A)** Distribution of the Kacac protein based on the site number per protein. **(B)** The consensus sequence logos show enrichment of amino acid residues among the Kacac sites. **(C)** Venn diagram shows the cellular compartment distribution of Kacac proteins. **(D)** Representative ontology annotations and all KEGG pathways enriched within the Kacac proteome. **(E)** Two protein complexes significantly enriched in the Kacac proteome. The color bar depicts the number of Kacac sites identified in each protein.

To investigate the subcellular distribution of Kacac substrates within cells, we conducted a cellular compartment analysis of the Kacac proteome. Previous studies have indicated that Ksucc and Kmal are notably enriched in mitochondria, while the subcellular localization of Kac, Khib, and Kbhb substrates is often observed in either the cytoplasm or the nucleus (9,20,38,39,41). The subcellular distribution of Kacac closely resembles that of Kac, Khib, and Kbhb substrates, with only 17% Kacac proteins are located in mitochondrion. Also, Kacac proteins are significantly more represented in the nucleus, accounting for 85% of all proteins (Fig. 5C).

To discern the physiologically relevant Kacac sites, we compared our dataset with public databases of known PTMs in UniProt (http://www.uniprot.org). Through analysis, we discovered 73 Kacac sites overlapping with other modifications (Supplementary Table S2A), encompassing ubiquitination and previously reported Kac, butyrylation (Kbu), Kcr, propionylation (Kpr), lactylation (Kla), Kbhb, Ksucc, Kmal, glutarylation (Kglu), Khib, as well as lysine methylation (Kme) sites. These PTMs may compete for the same lysine residues and reciprocally influence one another. For example, p53 Kbhb competes with p53 Kac, leading to reduced cell growth arrest and apoptosis (21). In principle, Kacac could occupy one of the same residues targeted by Kbhb and Kac. Moreover, we conducted a comparison between the Kacac sites and reported mutations linked to protein biological functions in the UniProt database. The findings revealed seven lysine sites significantly impacted by mutations, leading to altered protein functions (Supplementary Table S2A). Protein methylation and acetylation play pivotal roles in mediating crucial biological processes and signaling pathways. In our findings, several crucial proteins known as substrates for acetylation or methylation were discovered to be acetoacetylated, including p53, histone H2A.Z, and SMC3. Acetoacetylation at K370 of p53 may contribute to the regulation of p53’s activation or repression, thereby influencing rapid responses to DNA damage or cellular stress (42). Similarly, Kacac at K4 of histone H2A.Z may impaired GCN5 mediated H2A.Z.1 acetylation, thereby affecting RNA polymerase II recruitment and gene transactivation (43). In addition to methylation and acetylation, Kacac might influence the interaction of regulatory proteins with their protein partners by interfering with other reversible post-translational modifications (PTMs), such as the small ubiquitin-like modifier (SUMO) conjugation pathway. One such example is the identification of acetoacetylation at K102 of SERBP1. However, SUMOylation of SERBP1 at K102, K228, and/or K281 by small ubiquitin-like modifier (SUMO) proteins is implicated in the regulation of As_2_O_3_-induced PML nuclear bodies (PML-NBs) formation, which is associated with oxidative stress and interferon stimulation (44). Furthermore, four lysine residues (K38KKK) located in the N-terminal domain of caspase-7 facilitate the rapid proteolysis of poly(ADP-ribose) polymerase 1 (PARP-1), ensuring swift cell demise during apoptosis (45). Therefore, acetoacetylation of K38 (K38acac) may reduce the ability of caspase-7 to cleave PARP1. These findings suggest a potential interplay between Kacac and other PTMs within protein contexts, offering insights into the biological functions of Kacac and its role in complex metabolic regulation. In addition to competing with other modifications for the same residues, we identified 78 Kacac sites located within five residues of various types of PTMs, including lysine acylation, phosphorylation, O-GlcNAcylation, ubiquitination, and ADP-ribosylation. Notably, some of these Kacac sites are situated near critical mutation sites associated with protein dysfunction (Supplementary Table S2B). Together, these results imply the widespread and profound biological impacts of Kacac by functionally influencing substrates across the proteome.

### Functional annotation of the lysine acetoacetylome

To elucidate the biological functions of Kacac substrates in mammalian cells, we conducted a gene ontology (GO) analysis. Our findings revealed significant enrichment of Lys-acetoacetylated proteins in cellular metabolic processes, particularly in nucleosome assembly (p = 2.82 × 10^−13^) and gene expression (p = 3.98 × 10^−5^). Nucleosome assembly and disassembly are essential aspects of chromatin dynamics, critically involved in DNA replication, transcription, and repair. During nucleosome assembly and disassembly, histone chaperones bind to histones to prevent their abnormal aggregation and improper interactions with DNA, thereby facilitating their proper transfer onto DNA chains to form nucleosomes (46). Interestingly, we identified several histone chaperones that can undergo lysine acetoacetylation, including SET protein, NPM, and SSRP1, a component of the FACT (facilitates chromatin transcription) complex. In addition, ATP-dependent chromatin remodeling complexes modulate chromatin packing by sliding, ejecting, or restructuring nucleosomes, thereby enabling dynamic regulation of chromatin architecture (47). We also identified Kacac-modified substrates within ATP-dependent chromatin remodeling complexes, including BAZ1A from the ISWI complex and SMARCC2 from the SWI/SNF complex. These findings highlight the potential significant role of Kacac in nucleosome assembly.

Of further interest, we found that the molecular functions attributed to Kacac substrates included structural constituent of chromatin (p = 1.41 × 10^−17^), nucleosomal DNA binding (p = 2.75 × 10^−14^), protein heterodimerization activity (p = 1.26 × 10^−8^), histone binding (p = 1.92 × 10^−5^), and RNA polymerase II-specific DNA-binding transcription factor binding (p = 5.39 × 10^−5^) (Fig. 5D). The Kyoto Encyclopedia of Genes and Genomes (KEGG) pathway enrichment analysis indicated that Kacac proteins were enriched in immune response, RNA/ DNA metabolism, amino acid metabolism and cell cycle (Fig. 5D). Notably, most Kacac proteins are involved in systemic lupus erythematosus (p = 2.49 × 10^−21^), RNA degradation (p = 1.45 × 10^−2^) and mismatch repair (p = 2.59 × 10^−2^). To characterize the classification of the Kacac proteome, we conducted a protein class enrichment analysis. Chromatin/chromatin-binding, or -regulatory protein, HMG box transcription factor and RNA metabolism protein were prominently represented (Supplementary Fig. S6A). Chromatin-binding or regulatory proteins and HMG box transcription factors play key roles in regulating chromatin architecture, gene expression, DNA repair, and other cellular processes. In addition to the proteins identified in these two chromatin-related classes, a substantial number of Kacac-modified proteins are associated with RNA processing. For example, splicing factor 3b subunit 1 (SF3B1) is a core component of the U2 snRNP at the spliceosome’s catalytic center, where it is essential for recognizing and defining the 3′ splice site at intron–exon junctions (48). Other proteins, including HNRNPL and HNRNPC from the heterogeneous nuclear ribonucleoprotein (hnRNP) family, as well as NONO (non-POU domain-containing octamer-binding protein), also play crucial roles in RNA splicing (49,50). Collectively, these findings suggest potential regulatory roles for Kacac in chromatin remodeling, gene expression, and RNA processing.

To identify protein complexes regulated by Kacac, we performed protein complex enrichment analysis with the manually curated CORUM database (51). Our analysis revealed significant enrichment of Kacac substrates in several protein complexes (Supplementary Table S3), including the Nop56p-associated pre-rRNA complex (p = 7.46 × 10^−9^), the H2A.Z histone-exchange complex p400/TIP60 (p = 7.98 × 10^−9^), the LARC complex (LCR-associated remodeling complex) (p = 6.01 × 10^−8^), the TLE1 corepressor complex (p = 6.97 × 10^−8^), the MASH1 promoter-coactivator complex (p = 1.25 × 10^−7^), the SNF2h-cohesin-NuRD complex (p = 1.73 × 10^−6^), the PID complex (p = 1.81 × 10^−6^) and the Spliceosome, E complex (p = 8.85 × 10^−5^) (Fig. 5E, Supplementary Table S3). The Nop56p-associated pre-rRNA complex is linked to ribosome biogenesis in human cells (52). The H2A.Z histone-exchange complex p400/TIP60 regulates H2A.Z deposition, thereby influencing chromatin structure and gene expression (53). Our data revealed that the highest number of Kacac-modified substrates belong to these two complexes, with acetoacetylation detected in 9 of 104 subunits of the Nop56p-associated pre-rRNA complex and 6 of 27 subunits of the H2A.Z histone-exchange complex p400/TIP60, respectively. Notably, several components of the Nop56p-associated pre-rRNA complex were extensively modified with Kacac, including NUCL, NPM, and NOLC1 (Fig. 5E). These findings highlight the potential involvement of Kacac in protein complexes associated with chromatin remodeling, gene expression, and ribosome biogenesis. Additionally, the LCR-associated remodeling complex (LARC) exhibits a sequence-specific binding to the hypersensitive 2 (HS2)-Maf recognition element (MARE) DNA and orchestrates nucleosome remodeling (54). Spliceosomes catalyze the splicing of primary gene transcripts (pre-mRNA) to produce messenger RNA (mRNA). Our findings reveal that 5 out of 19 subunits within the LARC complex and 6 out of 129 subunits in the Spliceosome E complex undergo acetoacetylation, affirming the significance of Kacac in nucleosome assembly and a correlation between Kacac and RNA processing. Moreover, the TLE1 corepressor complex is recognized for its involvement in HES1-mediated transcriptional repression (55). Intriguingly, our analysis revealed acetoacetylation in 4 of the 8 subunits of this complex, with NUCL presenting 10 Kacac sites and the crucial protein PARP1 showing 2 Kacac sites. The HDAC1-containing PID complex, which is involved in repressing p53 transactivation functions and contributes to the regulation of p53-mediated growth arrest and apoptosis (56), is also enriched in our study. Notably, our data revealed acetoacetylation in 3 out of 5 subunits within the PID complex, shedding light on the contribution of Kacac to cell growth and proliferation. Together, these findings suggest that Kacac participates in various cellular functions, including nucleosome remodeling, RNA metabolism, and transcriptional regulation.

### Profiling physiological relevance of Kacac modification

To explore the epigenetic role of Kacac in gene regulation, we performed RNA sequencing (RNA-seq) from control and acetoacetate-treated HEK293T cells. We compared the differentially regulated genes between the two conditions and conducted GO analysis, KEGG pathway enrichment analysis, and gene set enrichment analysis (GSEA) to further understand the biological implications of Kacac. In our analysis, 4764 genes were found to be upregulated, whereas 2822 genes were downregulated after acetoacetate treatment (|fold change| > 1.5, adjusted p value < 0.05) (Fig. 6A, Supplementary Tables S4A, and S4B). We extracted differentially expressed genes (DEGs) between the "control" and "AcAc" conditions and generated a heatmap illustrating the expression patterns of the DEGs across both conditions (Supplementary Fig. S6B). GO analyses revealed that genes primarily enriched in downregulated terms are associated with metabolic processes such as amino acid metabolic processes, carboxylic acid catabolic processes, fatty acid catabolic processes, and pteridine-containing compound metabolic processes (Fig. 6B, Supplementary Table S5A). Additionally, DNA/RNA-associated processes such as tRNA metabolic processes, pyrimidine nucleotide metabolic processes, DNA replication, and DNA repair are prominently represented among downregulated terms (Fig. 6B). Several other processes, such as mitochondrial gene expression, translation, cell cycle checkpoint signaling, and protein localization to peroxisomes, are substantially downregulated following acetoacetate treatment (Supplementary Table S5A). In contrast, upregulated DEGs are predominantly enriched in processes including muscle contraction, cell adhesion, cellular component organization, regulation of nervous system processes, second-messenger-mediated signaling, regulated exocytosis, and leukocyte migration (Fig. 6B). Moreover, peptidyl-tyrosine phosphorylation, regulation of blood circulation, heart processes, and the phospholipase C-activating G protein-coupled receptor signaling pathway exhibit notably elevated expression following lithium acetoacetate treatment (Supplementary Table S5B). In line with the GO analysis findings, subsequent KEGG pathway enrichment analysis revealed that downregulated DEGs induced by acetoacetate are significantly enriched in pathways related to peroxisome function, cofactor biosynthesis, DNA replication and repair, amino acids and carbon metabolism (Fig. 6C, Supplementary Table S6A). Conversely, upregulated DEGs are predominantly associated with pathways such as neuroactive ligand-receptor interaction, the MAPK signaling pathway, the calcium signaling pathway, systemic lupus erythematosus, cytokine-cytokine receptor interaction, among others (Fig. 6C, Supplementary Table S6B). Additional GSEA results revealed enrichment of genes related to allograft rejection, KRAS signaling, muscle differentiation and inflammatory response (Fig. 6D). These findings further support that acetoacetate-mediated Kacac is closely linked to cellular processes involving amino acid metabolism, DNA/RNA metabolism, gene expression, translation, proliferation, and immune response.

**Fig. 6.**
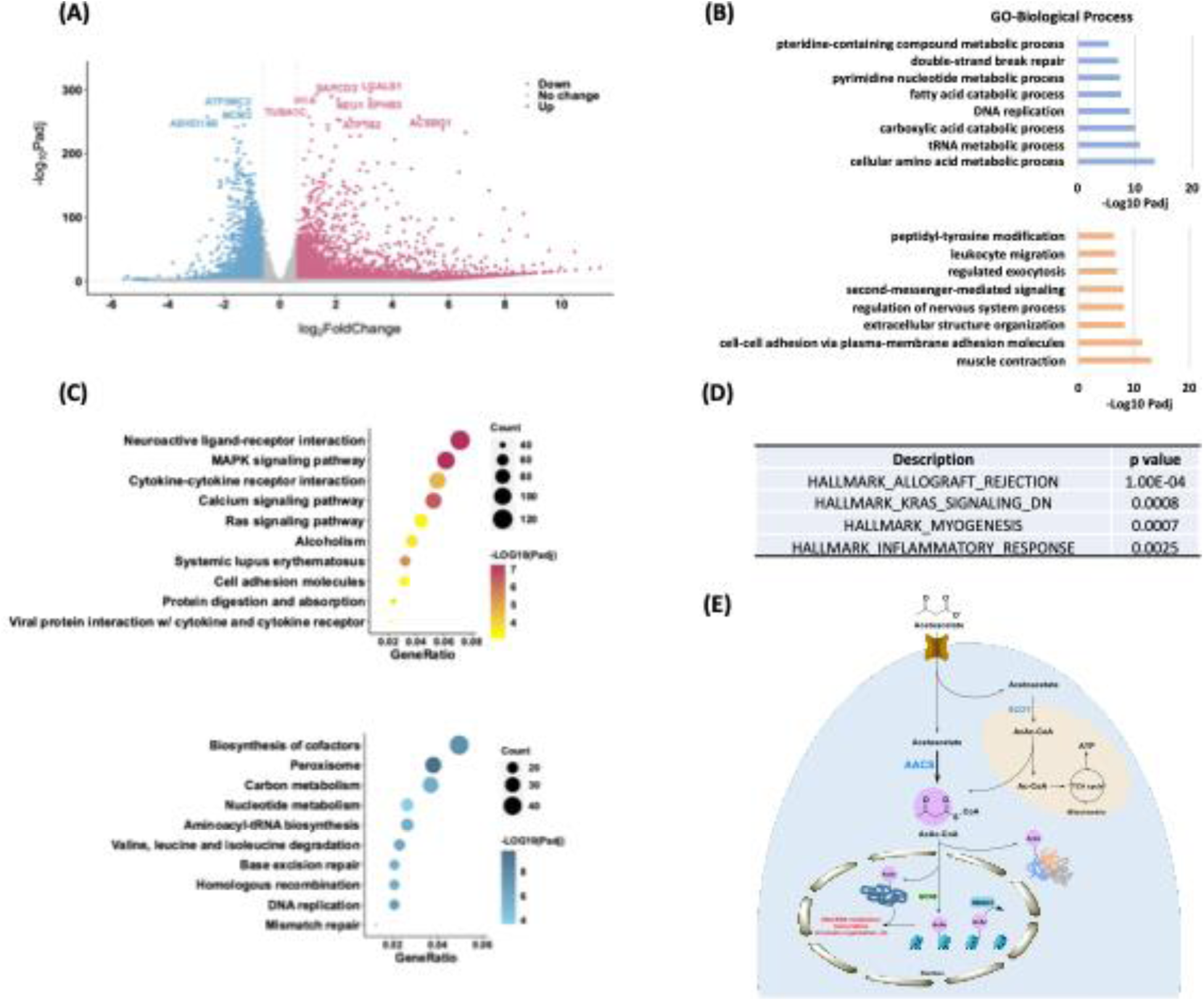
Profiling of physiological relevance of Kacac mark in HEK293T cells. **(A)** Volcano plot analysis of pairwise comparison of RNA-seq results from HEK293T cells with or without 20 mM acetoacetate treatment. **(B)** GO (Biological process) enrichment analysis of downregulated (blue) and upregulated (orange) differentially expressed genes (DEGs) after lithium acetoacetate treatment in HEK293T cells, ranked on the basis of adjusted p values. **(C)** Bubble plots showing the top 10 KEGG pathways enriched in the upregulated (upper) and downregulated (bottom) DEGs after lithium acetoacetate treatment in HEK293T cells, ranked on the basis of adjusted p values and counts. Gradient colors represent enriched significance, and size of circles represents numbers of DEGs. **(D)** Hallmark gene sets identified by GSEA after lithium acetoacetate treatment in HEK293T cells, ranked on the basis of P values. **(E)** A graphical model of Kacac. In this model, AACS, not SCOT, is a major player for AcAc-CoA and Kacac generation from acetoacetate.

## Discussion

Intracellular short-chain fatty acids and their corresponding acyl-CoAs play a pivotal role in regulating short-chain lysine acylations on cellular proteins, establishing a link between cellular metabolism and gene expression (6,23). Ketone bodies are crucial metabolites in maintaining physiological homeostasis and vital metabolic and signaling mediators in diverse physiological and pathological states (13). Recent studies have reported that β-hydroxybutyrate serves as a precursor for Kbhb on both histones and non-histones, significantly expanding its role as a protein function modifier (7,20). The Kbhb levels are dynamic in response to altered physiological conditions such as starvation and diabetic ketoacidosis (7).

Despite the recent disclosure unveiling histone Kacac as a novel PTM (22), the mechanisms underlying the dynamics and proteome-wide distribution for Kacac marks are largely unexplored. A thorough elaboration of the regulatory elements governing Kacac substrate proteins, along with the enzymes responsible for adding and removing this modification, is crucial for the functional characterization of Kacac and understanding the role of acetoacetate in cellular regulation. In this study, we developed a chemo-immunological approach for Kacac detection and applied it to conduct the first global proteomic analysis of the lysine acetoacetylome in human cells. The approach not only offers a reliable and time-efficient method for detecting Kacac without the need to develop new antibodies but also importantly enables the simultaneous detection and quantification of Kbhb and Kacac in the same biological samples. In our proteomic profiling, we identified 139 lysine acetoacetylated peptides across 85 proteins in HEK293T cells. We found that Kacac extensively modifies numerous biologically important proteins, notably at critical sites, indicating its significant roles in diverse cellular processes and disease development. Notably, we discovered 14 previously unidentified histone Kacac marks such as H3K4, H2BK5, H2BK20, H2BK108, H2AK9, and several marks on H1 and H2A.Z. Many of the identified Kacac sites overlapped with, or were in close proximity to, other PTMs, including acylation, methylation, phosphorylation, O-GlcNAcylation, ubiquitination, and ADP-ribosylation. The crosstalk between Kacac and other PTMs may influence protein localization, interactions with binding partners, or other functions, ultimately leading to altered biological outcomes. For example, acetoacetylation of NOLC1 at K579—located within the CK2α binding region (residues 568–596) and near Ser574, which significantly enhances CK2α binding affinity—may influence phosphorylation at Ser574 and NOLC1’s interaction with CK2α, potentially altering CK2’s phosphorylation activity and its downstream signaling events (57). Furthermore, we found that Kacac proteins exhibited a notable enrichment in the nucleus. In line with this finding, Kacac substrates were concentrated in proteins associated with various nuclear biological processes and signaling pathways, including chromatin organization, DNA repair, RNA metabolism as well as gene expression. Moreover, a considerable fraction of Kacac proteins was identified in the cytosol, while only a minor proportion was observed in mitochondria, mirroring the subcellular distribution patterns of Kac, Khib, and Kbhb substrates (20,38,39). However, this distribution differs from that of Ksucc or Kmal substrate proteins (9,41), which are predominantly localized in mitochondria, indicating the dynamic functions of different lysine acylations.

Structurally different lysine acylations correlate with distinct physiological states and may lead to functional consequences. Sequence preference analysis revealed an enrichment of both negatively charged amino acids and positively charged lysine, with proline being largely depleted at most positions related to the Kcr or Khib sites (37,39). Near the Ksucc sites, glycine, alanine, and lysine were preferentially located, whereas arginine and serine were predominantly depleted at most positions (9). While sharing most sequence preference similarities with the reported Kbhb (20), the flanking sequence analysis of Kacac substrates revealed a higher enrichment of alanine at more positions near Kacac sites. Functional analysis further substantiates the unique and distinct regulatory roles of various acylations. For instance, Kac predominantly targets proteins involved in RNA biology, while Ksucc and Khib are notably enriched in multiple metabolism-related pathways (9,38). As the two primary ketone bodies, acetoacetate-mediated Kacac does indeed exhibit similarities to β-hydroxybutyrate-mediated Kbhb in terms of functional annotation, both of which are associated with DNA repair, RNA metabolism, and chromatin organization. However, Kbhb and Kacac still play distinct roles in specific processes. For instance, most Kbhb-modified proteins are involved in spliceosome, ribosome function and RNA transport, whereas Kacac-modified proteins are enriched in systemic lupus erythematosus and RNA degradation pathways. Compared to previously reported acylated substrates, 43 out of 85 Kacac proteins are preferred substrates of various acylation marks, including Kbz, Khib, Kac, and Kbhb (20,38,39,58). These overlapping proteins participate in key cellular processes such as chromatin remodeling, gene expression, apoptosis, and DNA/RNA metabolism—including DNA repair, RNA biosynthesis, and RNA splicing—highlighting the shared functional roles of protein lysine acylations. Among the 139 Kacac sites, 69, 44, 47, and 84 overlapped with Kbz, Kac, Khib, and Kbhb sites, respectively, with Kbhb showing the highest degree of overlap with Kacac compared to the other PTMs. Additionally, comparison of the Kacac proteome with the reported Kbhb-modified proteins reveals that 75 out of 85 proteins carry both modifications, indicating significant overlap in substrate specificity between these two ketone body–mediated PTMs. Although different PTM markers can occupy the same lysine sites on overlapping substrate proteins, their functional consequences might be distinct. For example, PARP1 is activated by DNA damage and catalyzes the ADP-ribosylation of multiple proteins, leading to the recruitment of DNA repair factors and chromatin remodeling (59). Interestingly, lactylation of PARP1 at seven lysine sites (K498, K505, K506, K508, K518, K521 and K524) enhances its ADP-ribosylation activity, whereas acetylation reduces it, indicating that lysine acylations differentially regulate PARP1 activity and may influence the DNA damage response (60). Furthermore, some Kacac sites coincide with residues previously reported to be modified by other lysine acylations implicated in disease progression. For example, malonylation of NUCL at K124 and K398 triggers its translocation and promotes AKT translation, thereby driving cell proliferation and tumor growth in hepatocellular carcinoma (HCC) (61). Acetylated NPM at K229 and K230 functions as a coactivator during RNA polymerase II-driven transcription and regulates the expression of genes that promote oral tumorigenesis (62). Another study reported that lactylation of chromobox 3 (CBX3) at K10 promotes its interaction with H3K9me3 and facilitates gastrointestinal cancer progression (63). These findings suggest a potential interplay between Kacac and other lysine acylation marks at protein sites, as well as a pathological significance of Kacac, although its precise functions remain to be elucidated.

In a prior study, p300 was identified as a β-hydroxybutyryltransferase, while HDAC1 and HDAC2 were characterized as de-β-hydroxybutyrylases (20). In our biochemical screening, we demonstrated that p300 and GCN5 function as acetoacetyltransferases. This finding is consistent with a previous study that highlighted the modest activity of p300 and substantial activity of GCN5 in transferring the acetoacetyl motif onto histone proteins (22). p300 has been shown to catalyze a wide range of acylations—including propionylation, butyrylation, crotonylation, 2-hydroxyisobutyrylation, succinylation, and β-hydroxybutyrylation—owing to its unique acyl-CoA binding pocket (6,29,64–66). The absence of such a pocket results in the limited activity of GCN5, which efficiently catalyzes propionylation but is poorly active for butyrylation and crotonylation (67,68). Surprisingly, GCN5 appears to exhibit relatively higher acetoacetyltransferase activity compared to p300; however, further evidence is needed to clarify their catalytic mechanisms as acetoacetyltransferases. We found that PCAF also exhibits robust lysine acetoacetyltransferase activity. Given that PCAF possesses conserved functional domains and exhibits high sequence similarity with GCN5 (75% amino acid identity) (69), it is not surprising that our data highlight its notable activities in vitro. HBO1 is a versatile protein acyltransferase that catalyzes acetylation, propionylation, butyrylation, crotonylation, lactylation, benzoylation, and acetoacetylation (22,58,70,71). A previous study elucidated that the CoA moiety, rather than the acyl group of acyl-CoAs, plays a major role in binding to HBO1 (70). Unfortunately, our test did not demonstrate the acetoacetyltransferase activity of HBO1 as reported in a previous study (22). This may be attributed to the low activity of the recombinant HBO1 used in our assay. HBO1 has been reported to assemble into a multisubunit complex that includes ING4/5, hEaf6, and scaffold proteins from either the JADE1/2/3 or BRPF1/2/3 families, which strongly enhance its acyltransferase activity (70). Further investigation of the enzymatic kinetics of p300, PCAF, GCN5, and possibly other HATs, both in isolation and within their complexes, would provide valuable insights.

PCAF and GCN5 are involved in key processes through their roles as acetyltransferases. PCAF, for instance, acetylates p53, influencing the protein’s DNA-binding ability, and apoptotic activities following DNA damage (72,73). PCAF also induces acetylation of HMGA1 proteins in PC-3 human prostate cancer cells and acetylates HMGB1, which in turn regulates the release of HMGB1 and is linked to the cellular inflammatory response (74,75). Interestingly, we found that several substrates previously identified as acetylation targets of GCN5/PCAF, including p53 and high mobility group (HMG) proteins, can also undergo acetoacetylation. Using recombinant histone proteins as substrates, we demonstrated that acetoacetyl-CoA can compete with acetyl-CoA for binding to PCAF/GCN5. Therefore, it is of great interest to further investigate how GCN5/PCAF regulate dynamic PTMs, including Kacac, and their effects across various metabolic processes or pathological conditions. Beyond establishing p300, GCN5 and PCAF as Kacac writers, we demonstrate that HDAC3 acts as an eraser of Kacac *in cellulo*. Further research is needed to investigate the unique biological effects of HDAC3-mediated de-Kacac and to identify additional HDACs that possess de-Kacac activity under physiological conditions.

Short-chain acyl-CoAs, derived from their respective short-chain fatty acids, act as cofactors for acyltransferases, facilitating various lysine acylation reactions (7,23). Our data demonstrate that acetoacetate, rather than the catabolism of ketogenic amino acids leucine and lysine, primarily contributes to Kacac formation in histones. This observation is consistent with previous research showing that ketone bodies primarily originate from the oxidation of fatty acids, with leucine contributing only up to 4% of the carbon in ketone bodies (13,76). Herein, we investigated the mechanism by which acetoacetate generates acetoacetyl-CoA for histone Kacac. We explored the contributions of AACS and SCOT in utilizing acetoacetate to increase the acetoacetyl-CoA pool for histone Kacac. Interestingly, we found that AACS demonstrates a substantial capacity, while SCOT exhibits a lower ability, to convert exogenous acetoacetate for histone Kacac in HEK293T cells. Endogenous SCOT levels in HEK293T cells may be saturated for Kacac generation, explaining why SCOT overexpression did not significantly impact Kacac levels in our study, while SCOT inhibition did. These findings shed light on the mechanism by which AACS utilizes acetoacetate to generate acetoacetyl-CoA in cytosol, subsequently transferring it into the nucleus for histone Kacac. This notion is in line with an earlier report suggesting that nuclear acyl-CoAs can equilibrate with the cytosolic acyl-CoAs pool across nuclear pores, facilitating histone acylation and thereby influencing gene expression regulation (6,77). There is evidence supporting the existence of significant pools of CoA and acyl-CoAs in mitochondria and peroxisomes (78,79). However, it remains unclear whether and how acyl-CoAs synthesized in these compartments can contribute to the nuclear acyl-CoA pool for histone acylations. As a crucial mitochondrial ketolytic enzyme, SCOT may predominantly contribute to Kacac on mitochondrial proteins rather than nuclear histones. A recent study revealed that SCOT is widely distributed throughout the cell and serves as a lysine succinyltransferase for global lysine succinylation (Ksucc) on proteins in mammalian cells (25). Notably, SCOT mediates Ksucc on mitochondrial protein LACTB, leading to enhanced mitochondrial membrane potential and respiration in liver cancer cells (25). Therefore, it is also possible that SCOT is widely distributed throughout the cell and has additional roles in acetoacetate utilization beyond its function as a ketolytic enzyme in the mitochondria.

AACS expression greatly increased histone Kacac levels in HEK293T cells. In contrast, AACS did not show significant effects in increasing histone Kacac levels in HepG2 cells. It is possible that even if histone Kacac is regulated by AACS in HepG2 cells, its turnover rate is slow, or there may be limited availability of free CoA in the HepG2 cells, which could hinder the further generation of acetoacetyl-CoA after AACS overexpression. It is also possible that AACS regulation in HepG2 cells is more complex, and its activity is dynamically influenced by several factors. Given the low Km of AACS for acetoacetate and the consistent presence of acetoacetate at concentrations sufficient to saturate or nearly saturate AACS, the utilization of acetoacetate as an anabolic substrate is governed not by the level of acetoacetate production, but rather by the regulation of AACS expression and activity. The mRNA level or activity of hepatic AACS is intricately regulated by factors such as modulators of cholesterol synthesis, acyl-CoAs, and physiological states like fasting and obesity (24). Furthermore, the utilization of acetoacetate for lipid synthesis is significantly augmented in highly malignant hepatoma cells, as evidenced by its pathway remaining unaffected by hydroxycitrate inhibition (80). These findings support our hypothesis that AACS is intricately regulated by various factors in HepG2 cells, although additional evidence is needed to substantiate this claim. Ketone body metabolism through AACS plays a pivotal role in maintaining cholesterol homeostasis (81). Therefore, we further investigated the contributions of HMG-CoA reductase (HMGCR), a rate-limiting enzyme in the cholesterol biosynthetic pathway, to histone Kacac levels. However, neither the overexpression of HMGCR nor the blockade of cholesterol biosynthesis by lovastatin significantly influenced histone Kacac levels. This suggests that HMGCR-associated cholesterol biosynthesis does not compete for acetoacetate production, thus not affecting Kacac in HepG2 cells.

The ratio of acetoacetate to β-hydroxybutyrate is directly linked to the mitochondrial NAD^+^/NADH ratio, and the equilibrium constant of β-hydroxybutyrate dehydrogenase 1 (BDH1) favors the production of β-hydroxybutyrate (82,83). A cytoplasmic β-hydroxybutyrate dehydrogenase (BDH2), sharing 20% sequence identity with BDH1, possesses a high Km for ketone bodies and facilitates the exclusive NAD^+^-dependent conversion of cytosolic β-hydroxybutyrate to acetoacetate (84,85). This enzyme serves either as a secondary system for energy supply or as a contributor to generate precursors for lipid and sterol synthesis (84). BDH1 likely primarily facilitates the conversion between acetoacetate and β-hydroxybutyrate, which may explain our findings that acetoacetate induces Kbhb to some extent, while β-hydroxybutyrate treatment promotes Kbhb rather than Kacac. However, the extent of BDH1/BDH2 involvement and their regulatory role in the equilibration of acetoacetate and β-hydroxybutyrate for their corresponding PTMs requires further investigation.

The RNA-seq analyses revealed that acetoacetate extensively influences gene expression, particularly in processes/pathways involved in amino acid and carbon metabolism, DNA/RNA metabolism, DNA transcription, nervous system regulation, immune response, proliferation, and inflammatory response. These enriched processes/pathways closely overlap with those identified in our proteomic data, affirming the biological effects of acetoacetate-mediated Kacac. When comparing our RNA-seq data with previous data on Kbhb, we identified several pathways predominantly enriched in acetoacetate, rather than β-hydroxybutyrate treatment. These pathways include biosynthesis of cofactors, homologous recombination, the MAPK signaling pathway, alcoholism, and hypertrophic cardiomyopathy, indicating a distinctive regulatory pattern of acetoacetate. Interestingly, we noticed an inverse trend in the signaling pathways induced by acetoacetate compared to those induced by β-hydroxybutyrate. For instance, pathways such as peroxisome, valine, leucine, and isoleucine degradation, ribosome, and RNA degradation were suppressed following acetoacetate treatment, while they were upregulated in response to an increase in β-hydroxybutyrate levels (7). Conversely, pathways upregulated by acetoacetate are prominently enriched in neuroactive ligand-receptor interaction, the calcium signaling pathway, systemic lupus erythematosus, cytokine-cytokine receptor interaction, and cell adhesion molecules while these pathways are significantly downregulated in response to increased β-hydroxybutyrate levels (7). Similarly, disease-related pathways, including systemic lupus erythematosus and type I diabetes mellitus are upregulated by acetoacetate while significantly downregulated in response to β-hydroxybutyrate stimulation (7). Previous studies have shown evidence supporting that acetoacetate may possess an opposite repertoire of signaling functions compared to β-hydroxybutyrate in certain aspects. For instance, acetoacetate activates GPR43 and sustains energy homeostasis by regulating lipid metabolism under ketogenic conditions (86). However, β-hydroxybutyrate exerts an antilipolytic effect by activating the GPR109A receptor on adipocytes, thereby creating a negative feedback loop where ketosis suppresses lipolysis in adipocytes (87). Furthermore, high concentrations of acetoacetate may trigger a pro-inflammatory response whereas β-hydroxybutyrate exerts a predominantly anti-inflammatory response (13). Similarly, the administration of exogenous β-hydroxybutyrate induced hepatic fibrosis, while acetoacetate inhibited it (88). Collectively, these findings, along with our data, support the notion that acetoacetate may possess distinct functions and potentially form a balanced system with β-hydroxybutyrate to co-regulate cellular processes, partially achieved through Kacac and Kbhb.

With the discovery of Kacac as a new PTM in the proteome, many open questions surge to the biological and biomedical fields. The effectiveness of ketogenic diets in certain diseases is partly attributed to the SCOT-mediated terminal oxidation of acetoacetate. Evidence indicates a reduction in SCOT expression under diabetic conditions and during myocardial injury (89,90). Inborn errors in SCOT function or SCOT knockout lead to severe consequences such as ketoacidosis, lethargy, and even neonatal death (91,92). In addition to their role in mitochondrial function, ketone bodies may contribute to disease-related metabolic regulation by participating in *de novo* lipid synthesis. AACS, the key enzyme in ketone body utilization for lipid synthesis, is upregulated and implicated in the development and progression of hepatocellular carcinoma (28). Acetoacetate, rather than β-hydroxybutyrate, can be readily utilized via AACS and is the preferred substrate for lipid synthesis compared to glucose and acetate (93). Given the pivotal role of SCOT/AACS in critical regulatory effects and their potential mediation of Kacac, it is imperative to extensively explore the SCOT/AACS-mediated Kacac substrates to elucidate the mechanisms underlying acetoacetate-associated diseases. Although acetoacetate and β-hydroxybutyrate, as two main metabolic ketone bodies, are well recognized as catabolic substances, our proteomic data and RNA-seq data support that acetoacetate-mediated Kacac functions differently from β-hydroxybutyrate-mediated Kbhb and may form a balance cycle with Kbhb in certain pathways, adapting to (patho)physiological changes. Notably, acetoacetate displayed unique roles in cancer cells, including the promotion of tumor growth in BRAF-V600E+ cancer cells through the activation of MEK-ERK signaling and induction of FGF21 expression in HepG2 cells (76). All these studies underscore the importance of extensively investigating the unique regulatory functions of acetoacetate. Hence, further detailed functional studies of Kacac substrates in specific disease models can be conducted to gain a deeper understanding of the biological mechanisms of acetoacetate. Of further note, some protein readers (e.g. bromodomain, YEATS domain, and PHD domain) of PTM marks display varying binding affinities towards different PTM marks (6). Therefore, comprehending the distinct reader proteins that specifically recognize Kacac is crucial for investigating the diverse biological processes of Kacac that set it apart from other types of PTMs.

In summary, we innovated an enabling chemo-immunological approach for Kacac protein detection and enrichment, in combination with high resolution tandem mass spectrometer and functional analyses, to unveil acetoacetate-mediated Kacac as a pivotal molecular mechanism for the global modification of cellular proteins and the regulation of cellular metabolism. We discovered several regulatory elements of lysine acetoacetylation, expounding its dynamics in cells. Our study not only opens a new window for investigating the crosstalk between metabolism and epigenetic regulation but also lays a foundation for further exploring the diverse roles of Kacac in moderating environmental impacts on metabolism and diseases related to ketone bodies. Importantly, our study uncovers an additional avenue through the dynamic coordination of Kacac and Kbhb pathways in response to metabolic changes, contributing to the maintenance of cellular homeostasis.

## Materials and Methods Materials and reagents

Unless otherwise noted, all chemical reagents were purchased from Sigma-Aldrich (St. Louis, MO). For synthesis of the peptide, the following N^α^-Fmoc protected amino acids and reagents were purchased from ChemPep Inc. (Wellington, FL): Fmoc-Ser(tBu)-OH, Fmoc-Gly-OH, Fmoc-Lys(Boc)-OH, Fmoc-Ala-OH, Fmoc-Thr(tBu)-OH, Fmoc-Gln(Trt)-OH, Fmoc-Pro-OH, Fmoc-Val-OH, Fmoc-Asp(tBu)-OH, Fmoc-Glu(tBu)-OH, Rink Amide resin, and 1-Ethyl-3-(3-dimethylaminopropyl)carbodiimide hydrochloride (EDC HCl). Other chemicals used for peptide synthesis were N-methylpyrrolid-2-one (NMP), *p*-toluenesulfonic acid, and N-hydroxysuccinimide from Oakwood Chemical (Estill, SC); O-(1*H*-6-Chlorobenzotriazole-1-yl)-1,1,3,3-tetramethyluronium hexafluorophosphate (HCTU) and sodium chloride from Chem-Impex (Dale, IL); ethane dithiol and ethyl acetoacetate from Alfa Aesar (Haverhill, MA); thioanisole and trifluoroacetic acid (TFA) from Acros Organics (Geel, Belgium); diisopropylethylamine (DIPEA) from TCI America (Portland, OR); dimethyl sulfoxide (DMSO) and diethyl ether from Fisher Scientific (Hampton, NH); ethylene glycol and sodium hydroxide from VWR (Radnor, PA); sodium bicarbonate from Avantor (Radnor, PA); hydrochloric acid from Aqua solutions (Jasper, GA). Antibodies were the following: pan anti-Kbhb (PTM BioLabs, PTM-1201), pan anti-Kac (PTM BioLabs, PTM-105), anti-Flag (Thermo Fisher Scientific, Cat# MA1-91878), anti-myc (Thermo Fisher Scientific, Cat# MA1-21316), anti-β-actin (Santa Cruz Biotechnology, Cat# sc-47778), anti-H3 (Santa Cruz Biotechnology, Cat# sc-517576), goat anti-rabbit IgG-HRP antibody (Cell Signaling Technology, Cat# 7074S), anti-mouse (Cytek Biosciences, Cat# 72-8042). Recombinant histone H3 and H4 proteins were purchased from New England Biolabs. HepG2, HCT116 and HEK293T cell lines were purchased from American Type Culture Collection (ATCC) and used without further authentication.

## Chemical synthesis

Please find the synthesis of K15acac-H2B (1–26) peptide in the Supplementary Methods.

## Dot blot

The synthetic peptide was reduced with sodium borohydride (NaBH_4_) in sodium carbonate (Na_2_CO_3_) buffer (pH 9.0) at room temperature. The reaction mixture was then spotted on nitrocellulose (NC) membranes. After incubation with 5% non-fat milk for 1 h, the membrane was incubated with the anti-β-hydroxybutyryllysine antibody (Kbhb) overnight at 4 °C. After three washes with TBST (Tris-buffered saline, pH 7.4, 0.1% (v/v) Tween 20), the membrane was incubated with a 1:3000 dilution of goat anti-rabbit IgG-HRP antibody at room temperature for 1 h. Following an additional wash with TBST, chemiluminescent detection was performed using the ECL substrate (Thermo Fisher Scientific, Cat# 32209).

## Preparation of histones and cell lysate

HEK293T cells were cultured in DMEM (Corning, Cat# 10-013-CV) containing 10% FBS (Thermo Fisher Scientific, Cat# A5256801), 1% streptomycin-penicillin (Thermo Fisher Scientific, Cat# 15140122). Cells were treated with lithium acetoacetate (Fisher Scientific, Cat# A1478), sodium β-hydroxybutyrate, valine, or leucine followed by whole cellular proteome extraction or histones extraction. Cell lysates were extracted using M-PER™ Mammalian Protein Extraction Reagent (Thermo Fisher Scientific, Cat# 78501) and 1% Protease Inhibitor Cocktail III (Thermo Fisher Scientific, Cat# 78425) with gentle sonication. The remaining debris was removed by centrifugation at 13200 × rpm at 4 °C. Histone proteins were extracted with the EpiQuik Total Histone Extraction Kit (Epigentek, Cat# OP-0006-100) according to the manufacturer’s instructions. The protein concentration was determined by Bradford assay.

## NaBH_4_-Kbhb antibody method based western blot analysis

Protein extracts (20 µg of cell lysate or 7 µg of histones) were first reduced by NaBH_4_ in Na_2_CO_3_ buffer (pH 9.0) at room temperature. Samples after reduction were further fractionated by sodium dodecyl sulfate-polyacrylamide gel electrophoresis (SDS-PAGE) and transferred to a NC membrane. After incubation with 5% non-fat milk in TBST for 1 h at room temperature, the membrane was incubated with the anti-Kbhb antibody at 1:2000 dilution at 4 °C overnight. Then the membrane was washed three times with TBST and incubated with a 1:3000 dilution of goat anti-rabbit IgG-HRP antibody at room temperature for 1 h. Following three washes with TBST, the membrane was analyzed by chemiluminescence using the ECL substrate.

## Expression and purification of HATs

HAT1 (20-341), p300 (1287-1666) and MOF (125-458) were expressed and purified as described in the previous work of our lab (8,94). The expression and purification of MOZ protein (pET28a-LIC-MOZ plasmid, Addgene #25181) were done according to the protocol from Structural Genomics Consortium (SGC) (http://www.thesgc.org/structures/2ozu). Maltose binding protein (MBP)-MORF (361–716) was expressed and purified as described (95). Plasmids (pET28a-PCAF (493-658) and perceiver HBO1) were transformed into *Escherichia coli* BL21 (DE3)/RIL competent cells with heat-shock method followed by spreading cells on LB-Agar plate with kanamycin or ampicillin. Colonies were picked up and grown in 8 mL of Luria broth medium for 16 hours and then 1 L cultures of Luria broth medium containing kanamycin or ampicillin at 37 °C until the OD_595nm_ reach to 0.5-0.7. 0.3 mM of isopropyl β-d-1-thiogalactopyranoside (IPTG) was added to induce protein expression at 16°C overnight. Cells were collected by centrifugation and suspended in lysis buffer containing 50 mM sodium phosphate at pH 7.4, 0.25 M NaCl, 5% (v/v) glycerol, 0.1% (v/v) Triton X, 5 mM imidazole, 2 mM β-mercaptoethanol, and 1 mM Phenylmethylsulfonyl fluoride (PMSF). Cells were disrupted with a microfluidic cell disruptor followed by collection of supernatant. After equilibrating Nickel-NTA agarose resin with column buffer (20 mM Na-HEPES at pH 8.0, 300 mM NaCl, 10% (v/v) glycerol, 30 mM imidazole and 1 mM PMSF), the protein supernatant was loaded onto the resin. Next, the resin was washed with column buffer twice and washing buffer (20 mM Na-HEPES at pH 8.0, 300 mM NaCl, 10% (v/v) glycerol, 70 mM imidazole and 1 mM PMSF) for three times. The proteins on the resin were eluted with elution buffer (20 mM Tris-HCl at pH 8.0, 300 mM NaCl, 10% glycerol, 500 mM imidazole and 1 mM PMSF), and then dialyzed in the dialysis buffer (25 mM Na-HEPES/Tris-HCl at pH 8.0, 250 mM NaCl, 10% glycerol, 1 mM DTT) at 4°C overnight. The resultant proteins were concentrated by Millipore centrifugal filter. The protein concentration and purity were determined by using Bradford assay and SDS-PAGE, respectively. The proteins were then aliquoted and stored at −80°C.

## In vitro screening of HATs activities

1 μg of recombinant histone H3 or H4 were incubated with individual HAT enzymes and 50 μM acetyl-CoA or acetoacetyl-CoA in a HAT reaction buffer (50 mM HEPES-Na and 0.1 mM EDTA-Na, pH 8.0) at 30°C for 1 h. The reaction mixture was reduced by NaBH_4_, followed by the addition of SDS sample buffer. The levels of acetylation and acetoacetylation were assessed by western blot. To confirm the activities of HATs in vitro, synthetic histone peptides H3 (1–20) or H4 (1–20) (the sequence of H3 (1-20): Ac-ARTKQTARKSTGGKAPRKQL; the sequence of H4 (1–20): Ac-SGRGKGGKGLGKGGAKRHRK) were used as acyl acceptor substrates to allow enzymatic transfer of acetoacetyl-group by individual HAT in same reaction conditions. The reaction products were detected and validated by MALDI-MS.

## In vivo validation of HAT, HDAC3, AACS, SCOT and HMGCR activities

HEK293T cells or HepG2 cells were cultured and transfected with pCMVβ-p300-myc (Addgene, Cat# 30489), flag-HDAC3 (Addgene, Cat# 13819), AACS (OriGene Technologies, Cat# RC206247), SCOT (OriGene Technologies, Cat# RC203764) or HMGCR (Addgene, Cat# 86085) by using Lipofectamine 3000™ Transfection Reagent (ThermoFisher, Cat# L3000008). The cells were subsequently treated with 20 mM lithium acetoacetate for 24 hours, after which histones were extracted. Additional experiments were conducted by treating cells with acetohydroxamic acid or lovastatin in combination with lithium acetoacetate to study SCOT or HMGCR activity. Kacac levels were analyzed by western blot using the procedures aforementioned.

## Molecular docking

The referred proteins were imported into Maestro version 13.6.121 and subsequently prepared with filled in missing side chains in the preprocessing stage and optimized hydrogen bond assignments using PROPKA at pH 7.4. Imported acyl-CoAs were prepared with predetermined stereocenters with the generated states at pH 7.0 ± 2.0. The receptor grid was then generated by selecting key residues around the catalytic site and then specified by either maintaining key hydrogen bonding residues or through the use of a core constraint with the co-crystallized ligand. Lastly, ligand docking of the acyl-CoAs applied the constraints used in the receptor grid using extra precision sampling to yield the modeled poses of the acyl CoAs.

## Trypsin digestion of cell lysate

Extracted proteins were reduced with 50 mM LiBD_4_ in Na_2_CO_3_ buffer (pH 9.0) for 6 hours at room temperature. Whole proteome were precipitated by cold acetone overnight. Proteins were redissolved in 50 mM NH_4_HCO_3_ buffer (pH 8.0), followed by reduction with 20 mM DTT for 1.5 h at 37 °C, and alkylation with 40 mM iodoacetamide for 30 min at room temperature in darkness. Excess iodoacetamide was blocked by 10 mM DTT. Trypsin (Thermo Fisher Scientific, Cat# 90058) was added at 1:50 trypsin-to-protein mass ratio for protein digestion overnight at 37 °C. The enzymatic reaction was quenched by boiling the sample. Tryptic peptides were dried in a SpeedVac system (ThermoFisher Scientific).

## Peptide immunoprecipitation

Peptides were redissolved in NETN buffer (1 mM EDTA, 50 mM Tris-HCl, 100 mM NaCl, 0.5% NP-40, pH 8.0). Pan anti-Kbhb antibody was first conjugated to Protein A/G plus agarose (Santa Cruz Biotechnology Inc, Cat# sc-2003) and then incubated with tryptically digested peptides with gentle agitation overnight at 4 °C. The beads were then washed three times with NETN buffer, twice with ETN buffer (50 mM Tris-HCl pH 8.0, 100 mM NaCl, 1 mM EDTA) and twice with water. Peptides were eluted from the beads with 0.15% TFA and dried by SpeedVac system.

## HPLC-MS/MS analysis

The resulting peptides were dissolved in 0.1% formic acid in water and loaded onto a commercial C18 column Acclaim PepMap RSLCnano, 75 µm x 15 cm, 3 µm particle size (Thermo Fisher Scientific, Waltham, MA). The loaded peptides were separated using a gradient of 5% to 80% HPLC buffer B (0.1% formic acid in 80% acetonitrile, v/v) in buffer A (0.1% formic acid in water, v/v) at a flow rate of 300 nL/min over 180 min by Dionex Ultimate 3000 RSLCnano system (ThermoFisher Scientific, Waltham, MA). The samples were analyzed by an Eclipse™ tribrid orbitrap mass spectrometer (ThermoFisher Scientific, Waltham, MA). In positive ion mode, a 120 000 resolution full mass scan was collected, followed by data-dependent MS/MS using 28% higher collision energy (HCD) fragmentation at 30,000 resolution with cycle time of 3 sec. Charge state screening was enabled and precursors with a charge of +1, or an unknown charge were excluded. A dynamic exclusion duration of 60 s was enabled (96). A lock mass correction was also applied using a background ion (m/z 445.12002).

## Protein sequence database searching

The acquired MS/MS data was searched by Byonic (Protein Metrics; v4.1). All the data were searched against reviewed UniProt Human protein database (20,433 entries, http://www.uniprot.org) with decoy. Trypsin was specified as cleavage enzyme allowing a maximum of 2 missing cleavages. Cysteine carbamidomethylation was specified as fixed modification. Methionine oxidation, lysine acetylation, lysine methylation, lysine β-hydroxybutyrylation, deuterated lysine β-hydroxybutyrylation (DKbhb), lysine acetoacetylation, lysine propionylation and lysine butyrylation were included as variable modifications. FDR thresholds for protein, peptide and modification site were specified at 1%. Peptides with a Byonic PEP 2D value lower than 0.001, as well as those with PEP 2D values between 0.1 and 0.001 after manual confirmation, were retained for subsequent analysis.

## Bioinformatics analysis

Sequence preference motif was generated by iceLogo, utilizing the human proteome as the background (97). Mutations and recorded binding sites were extracted from UniProt database (http://www.uniprot.org). Gene ontology analysis was performed using PANTHER (version 18.0) (98). Enrichment analysis for KEGG pathway was carried out using GOstats package along with a hypergeometric test in R (99). Protein complex analysis was performed by using manually curated CORUM protein complex database for all mammals using a hypergeometric test (51). Protein complexes enriched in the Kacac proteome were visualized in Cytoscape (v3.10.1) (100).

## RNA-seq analysis

Total RNAs were extracted from control and lithium acetoacetate-treated (20 mM for 24 h) HEK293T cells using the RNeasy Plus Mini Kit (Qiagen, Cat# 74134). Three biological replicates were performed for each condition. Isolated RNA sample quality was assessed by High Sensitivity RNA Tapestation (Agilent Technologies Inc., California, USA) and quantified by AccuBlue® Broad Range RNA Quantitation assay (Biotium, California, USA). Paramagnetic beads coupled with oligo d(T)25 were combined with total RNA to isolate poly(A)+ transcripts based on NEBNext® Poly(A) mRNA Magnetic Isolation Module manual (New England BioLabs Inc., Massachusetts, USA). Prior to first strand synthesis, samples were randomly primed (5′ d(N6) 3′ [N=A,C,G,T]) and fragmented based on manufacturer’s recommendations. The first strand was synthesized with the Protoscript II Reverse Transcriptase with a longer extension period, approximately 30 minutes at 42 ⁰C. All remaining steps for library construction were used according to the NEBNext® Ultra™ II Directional RNA Library Prep Kit for Illumina® (New England BioLabs Inc., Massachusetts, USA). Final libraries quantity was assessed by Qubit 2.0 (ThermoFisher, Massachusetts, USA) and quality was assessed by TapeStation D1000 ScreenTape (Agilent Technologies Inc., California, USA). Final library size was about 450bp with an insert size of about 300bp. Illumina® 8-nt dual-indices were used. Equimolar pooling of libraries was performed based on QC values and sequenced on an Illumina® NovaSeq X Plus 10B (Illumina, California, USA) with a read length configuration of 150 PE for 40M PE reads per sample (20M in each direction). FastQC (version v0.12.1) was employed to check the quality of raw reads. Trimmomatic (version v0.39) was applied to cut adaptors and trim low-quality bases with default setting. STAR Aligner (version 2.7.10b) was used to align the reads. The package of Picard tools (version 3.0.0) was applied to mark duplicates of mapping. StringTie (version 2.2.1) was used to assemble the RNA-Seq alignments into potential transcripts. FeatureCounts (version 2.0.6) or HTSeq (version 2.0.3) was used to count mapped reads for genomic features such as genes, exons, promoter, gene bodies, genomic bins, and chromosomal locations. DESeq2 (version 1.40.2) was employed to process the differential gene expression analysis. The list of significance was established by setting the fold change threshold at a level of 1.5 and adjusted P < 0.05. The gene ontology (GO), gene set enrichment analysis (GSEA), and KEGG enrichment analysis were analyzed via the ClusterProfiler package and Molecular Signatures Database (MSigDB).

## Data Availability

All data are available in the main text or the supplementary materials. This study includes no data deposited in external repositories.

## Acknowledgements

We thank the Proteomics and Mass Spectrometry facility (PAMS) at UGA for the MS support. Proteomic analysis was performed at the Complex Carbohydrate Research Center and was supported in part by the National Institutes of Health (NIH)-funded R24 grant [R24GM137782 to Parastoo Azadi]. The Eclipse mass spectrometer used in the modified peptide analysis was supported by GlycoMIP, a National Science Foundation Materials Innovation Platform funded through Cooperative Agreement DMR-1933525. We are thankful to the National Institutes of Health [NIH 1R35GM149230] and the National Science Foundation [NSF 2203942] for grant support (PI Zheng). Author contributions: Q.F. and Y.G.Z. designed the research. Q.F. and T.N. performed experiments. Q.F., B.K. and P.A. conducted proteomic analysis. Q.F., T.N. and Y.G.Z. analyzed results. Y.G.Z. supervised the research. Q.F., T.N. and Y.G.Z. wrote the manuscript.

