## Supplementary File for "Identification of the Regulatory Elements and Protein Substrates of Lysine Acetoacetylation"

**This PDF file includes:**

Supplementary Methods
Figs. S1 to S6

**Other Supplementary Materials for this manuscript include the following:**

Tables S1 to S6

**Supplementary Methods**

Synthesis of K15acac-H2B (1-26) peptide

(1) 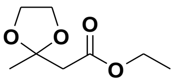 ethyl 2-(2-methyl-1,3-dioxolan-2-yl)acetate

To a flask, ethyl acetoacetate (1.96 mL, 15.368 mmol, 1 eq), ethylene glycol (1.12 mL, 19.978 mmol, 1.3 eq), and *p*-toluenesulfonic acid (29.2 mg, 0.154 mmol, 0.01 eq) were added to 20 mL of benzene, and refluxed overnight using a Dean-Stark trap. After cooling to room temperature, benzene was removed by rotary evaporation and diluted in 10 mL ethyl acetate. The organic solvent was then washed with 30 mL 5% aqueous sodium bicarbonate then 30 mL brine. The organic layer was dried over sodium sulfate before removal by rotary evaporation to obtain the product, ethyl 2-(2-methyl-1,3-dioxolan-2-yl)acetate as a colorless oil. Yield: 2.592 g, 96.82%. ESI-HRMS calc for $C_8H_{14}NaO_4$ $[M+Na]^+$: 197.0784 found 197.0779 $^1$H NMR (500 MHz, DMSO-$d_6$) δ 3.95 (q, $J$ = 7.1 Hz, 2H), 3.77 (s, 4H), 1.29 (s, 3H), 1.08 (t, $J$ = 7.1 Hz, 3H).

(2) 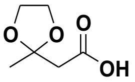 2-(2-methyl-1,3-dioxolan-2-yl)acetic acid

The ketal ester (1) (2.592 g, 14.88 mmol, 1 eq) was dissolved in 20 mL of ethanol before adding a 2M solution of sodium hydroxide (0.774 g, 9.67 mL, 19.344 mmol, 1.3 eq). The resulting solution was stirred at room temperature for 4 hours. The ethanol was removed by rotary evaporation and the resulting aqueous phase was subsequently washed with 2 x 20 mL of diethyl ether. The aqueous phase was then acidified with 2M HCl. The product was extracted from the aqueous phase with 3 x 100 mL of ethyl acetate. The organic phase was dried over sodium sulfate and then removed with rotary evaporation to obtain the product, 2-(2-methyl-1,3-dioxolan-2-yl)acetic acid as a yellow oil. 781 mg, 35.91%. ESI-HRMS calc for $C_6H_{10}NaO_4$ $[M+Na]^+$: 169.0471 found 169.0466 $^1$NMR $^1$H NMR (500 MHz, DMSO-$d_6$) δ 3.84 (s, 4H), 3.15 (s, 2H), 1.37 (s, 3H).

(3) 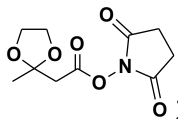 2,5-dioxopyrrolidin-1-yl 2-(2-methyl-1,3-dioxolan-2-yl)acetate

The protected acetoacetate (2) (598 mg, 4.09 mmol, 1 eq) was dissolved in 10 mL of dichloromethane before adding N-hydroxysuccinimide (753 mg, 6.54 mmol, 1.6 eq) and EDC HCl (1.25 g, 6.54 mmol, 1.6 eq) to the solution. The solution was stirred at room temperature for 3 hours. The solution was diluted with 100 mL of dichloromethane, and then the organic layer was washed with 50 mL of brine, 50 mL of saturated aqueous sodium bicarbonate, and finally 50 mL of brine. The organic layer was dried over sodium sulfate and subsequently evaporated to obtain the compound, 2,5-dioxopyrrolidin-1-yl 2-(2-methyl-1,3-dioxolan-2-yl)acetate as a yellow oil, with no further purification. 670 mg, 67.32%. ESI-HRMS calc for $C_{10}H_{13}NNaO_6$ $[M+Na]^+$: 266.0635 found 266.0628 $^1$H NMR (500 MHz, DMSO-$d_6$) δ 3.86 – 3.79 (m, 4H), 2.90 (s, 2H), 2.71 (s, 4H), 1.34 (s, 3H).

(4) Solid Phase Peptide Synthesis

The starting peptide (PEP AKS APA PKK GSK(Dde) KAV TKA QKK DG) was synthesized via standard Fmoc solid phase peptide synthesis using the FOCUS XC automatic synthesizer (AAPPTec, Louisville, KY). Rink amide resin was swollen in 15 mL of N,N-dimethylformamide (DMF) for 15 minutes and drained thereafter. 8 mL of 20% (v/v) piperidine in DMF was added and left to deprotect Fmoc for 10 minutes. This was repeated with a washing step before and after the second addition of piperidine. 5 mL of the corresponding protected amino acid (0.2 M in DMF, 5 equiv.) was activated in a separate vessel using 5 mL of HCTU (0.2 M in DMF, 5 equiv.) and 5 mL of 4-methylmorpholine (NMM) (0.2 M in DMF, 5 equiv.). The solution was mixed for 1 minute before being added to the resin. The reaction proceeded for 1 hour before proceeding to the draining and washing steps. The deprotection and coupling steps were repeated until the desired peptide sequence was acquired. Acetylation of the N-terminus was accomplished manually by adding a solution of 4:1 DMF to acetic anhydride (50 equiv.) and DIPEA (12.5 equiv.) to the resin. The reaction was allowed to mix for 30 minutes with subsequent draining and washing by DMF. Dde deprotection was accomplished manually as well by addition of 8 mL of 2% $N_2H_4$ in DMF. The reaction was mixed for 2 hours followed by the draining and washing of the resin. This was repeated once before additional washing and drying of the resin.

(5) Coupling acetoacetate

The peptide on resin was swollen in 20 mL DMF for 30 minutes then drained. A solution of the 5 mL of 2,5-dioxopyrrolidin-1-yl 2-(2-methyl-1,3-dioxolan-2-yl)acetate (0.2 M in DMF, 5 equiv.) and 5 mL of NMM (0.2 M in DMF, 5 equiv.) were added to the resin and mixed for 3 hours at room temperature. The resin was then drained, washed, and dried.

(6) Cleavage

Removal of the peptide from the resin was accomplished using 5mL of 95% TFA, 2.5% $H_2O$, and 2.5% triisopropylsilane (TIS) and allowed to mix for 4 hours at room temperature. The resulting solution was filtered to remove the resin and diluted with 40 mL of cold diethyl ether and centrifuged at 3214 g. The solution was decanted from the precipitate and 40 mL of cold diethyl ether was added again and followed by a second centrifugation step. The solution was decanted, and the precipitate was dissolved in water for high performance liquid chromatography (HPLC) purification using LC-20AT HPLC and SPD-20A UV/Vis detector (Shimadzu, Kyoto, Japan). The Agilent Eclipse XDB-C18 250 x 9.4 mm column (Agilent, Santa Clara, CA) was used for semi-preparative HPLC purification while the Agilent Eclipse XDB-C18 250 x 4.6 mm column (Agilent, Santa Clara, CA) was used for analytical HPLC to check peptide purity. The peptide was lyophilized to obtain a white powder. ESI-HRMS calc for $C_{122}H_{210}N_{36}O_{37}$ $[M]^+$: 2771.5657 found 2771.5587.

**(A)**

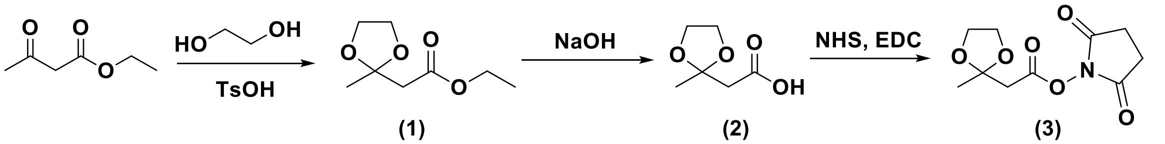


(1) Exact Mass: 174.08921

(2) Exact Mass: 146.05791

(3) Exact Mass: 243.07429

**(B)**

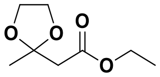

**ethyl 2-(2-methyl-1,3-dioxolan-2-yl)acetate**

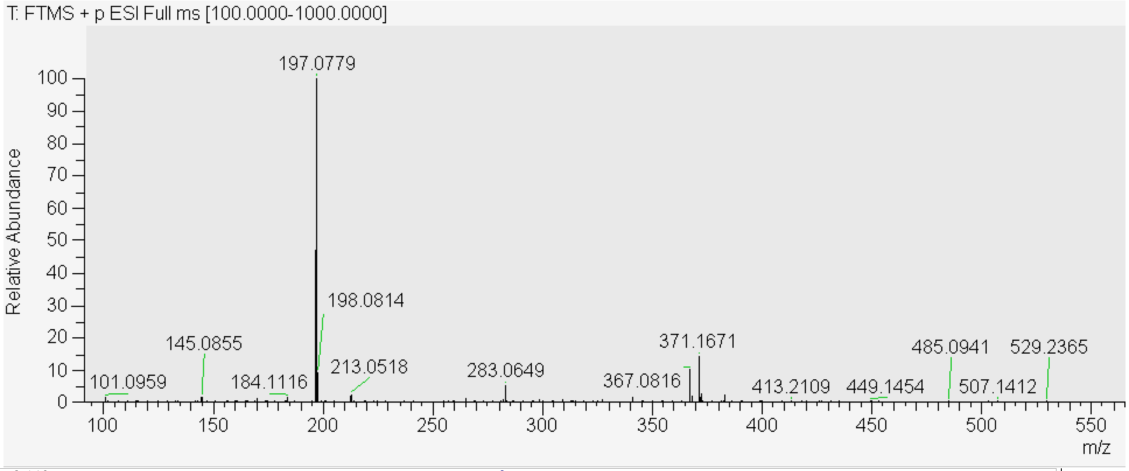


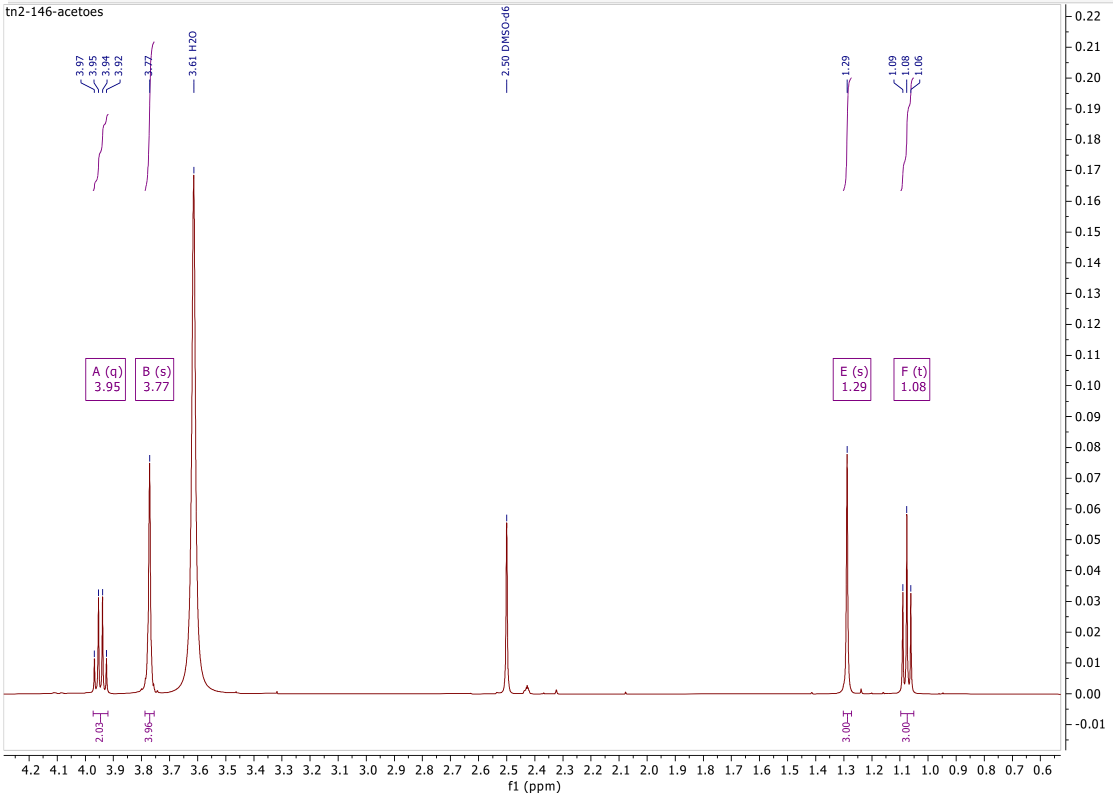

**(C)**

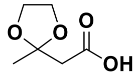


**2-(2-methyl-1,3-dioxolan-2-yl)acetic acid**

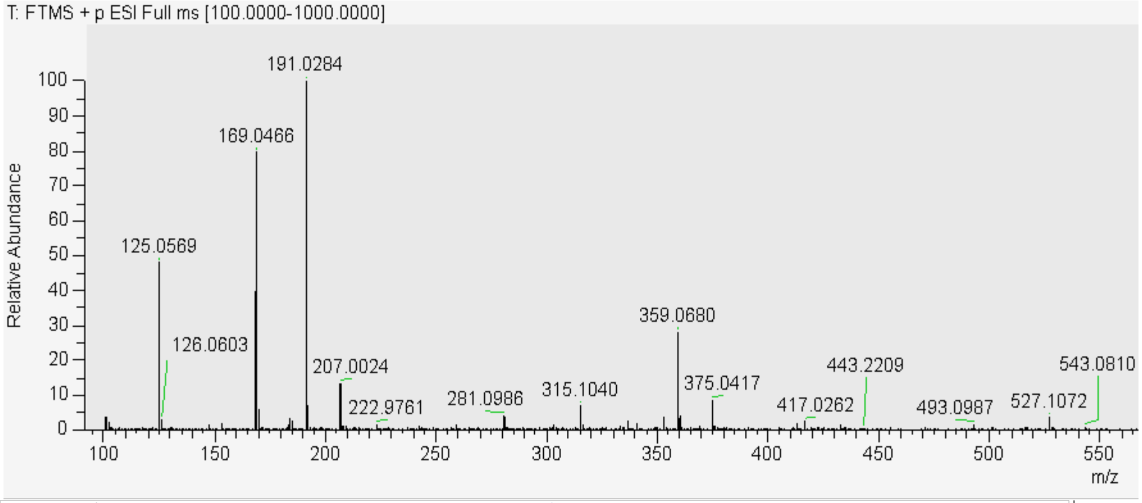


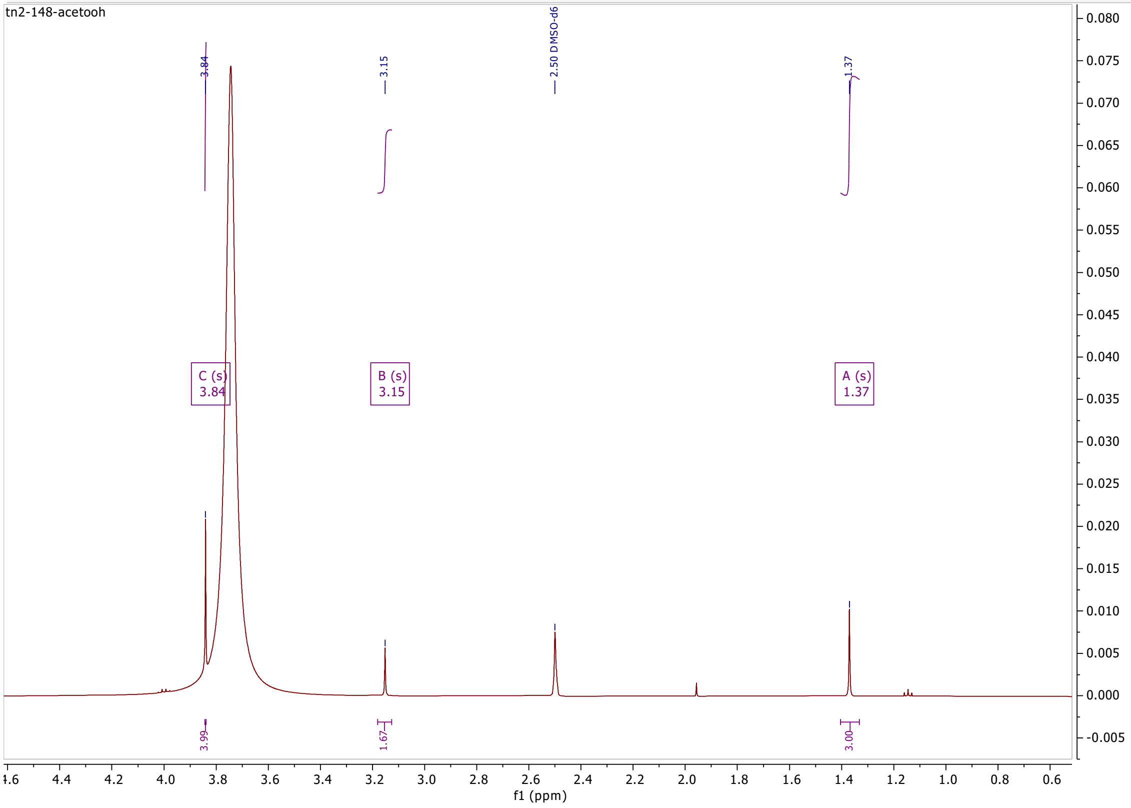

**(D)**

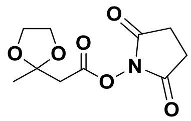


**2,5-dioxopyrrolidin-1-yl 2-(2-methyl-1,3-dioxolan-2-yl)acetate**

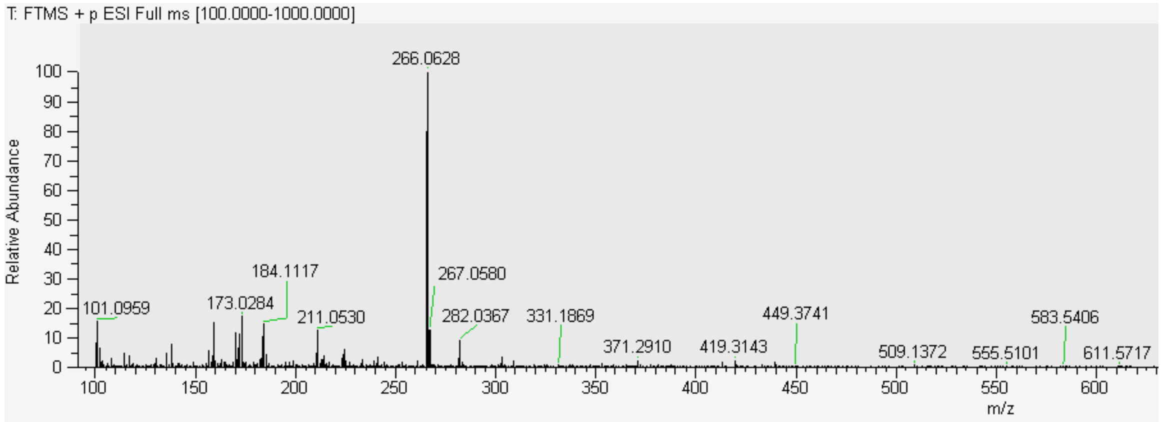


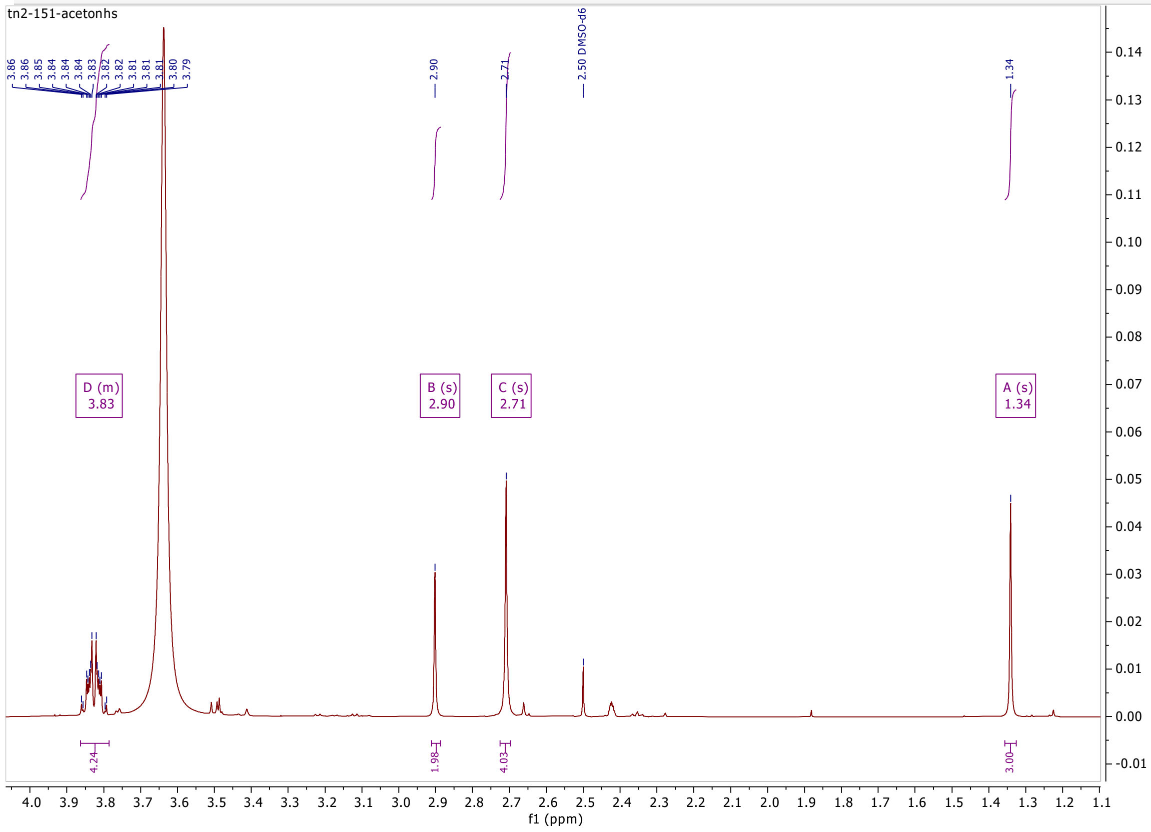

(E)

**Ac-PEPAKSAPAPKKGSKacacKAVTKAQKKDG-NH2**

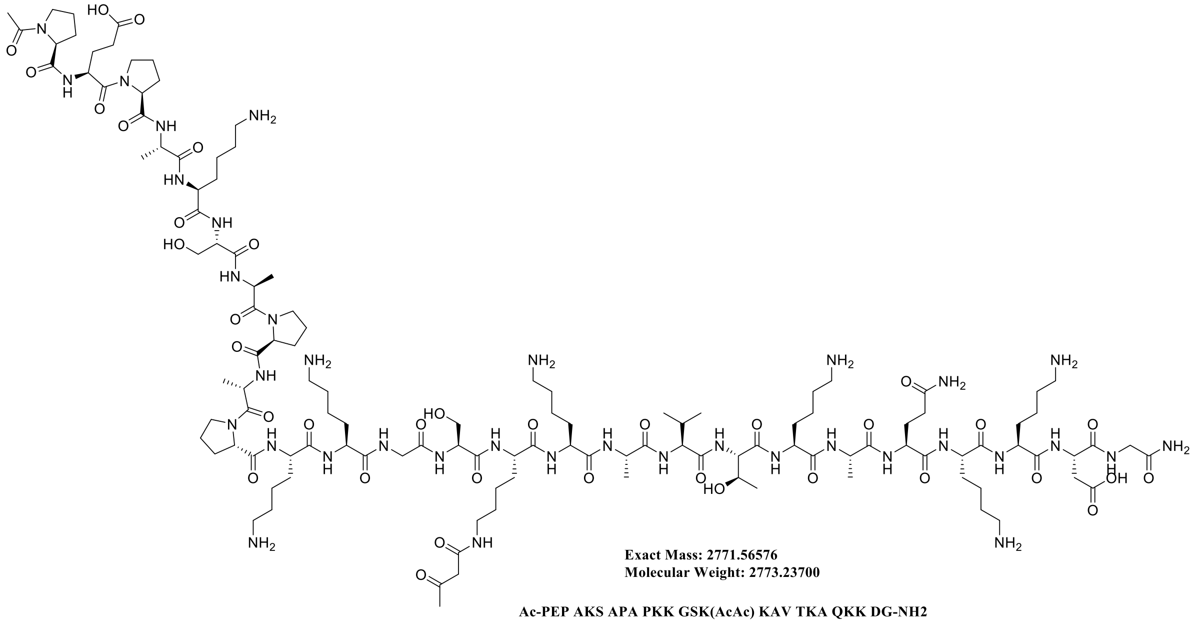


Ac-PEP AKS APA PKK GSK(AcAc) KAV TKA QKK DG-NH2

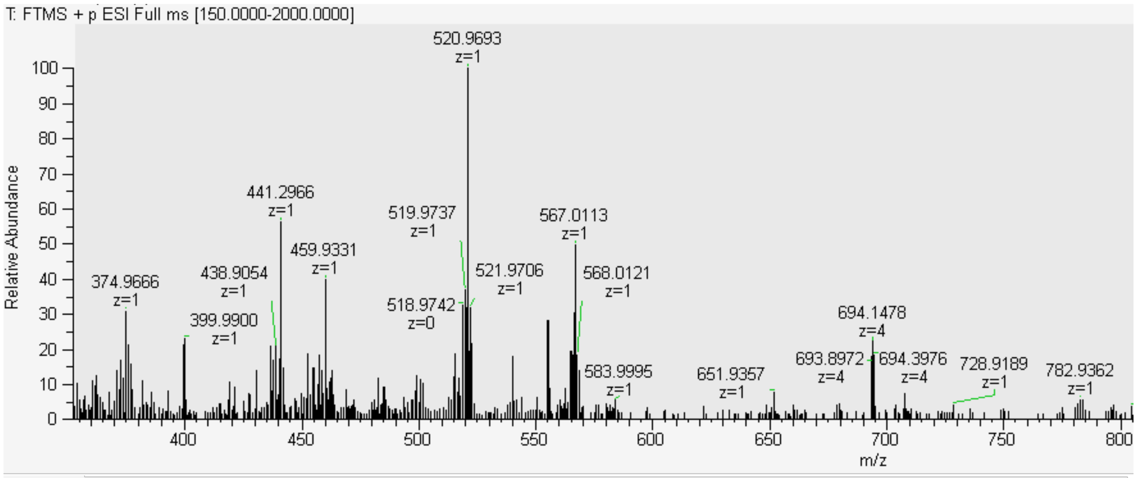


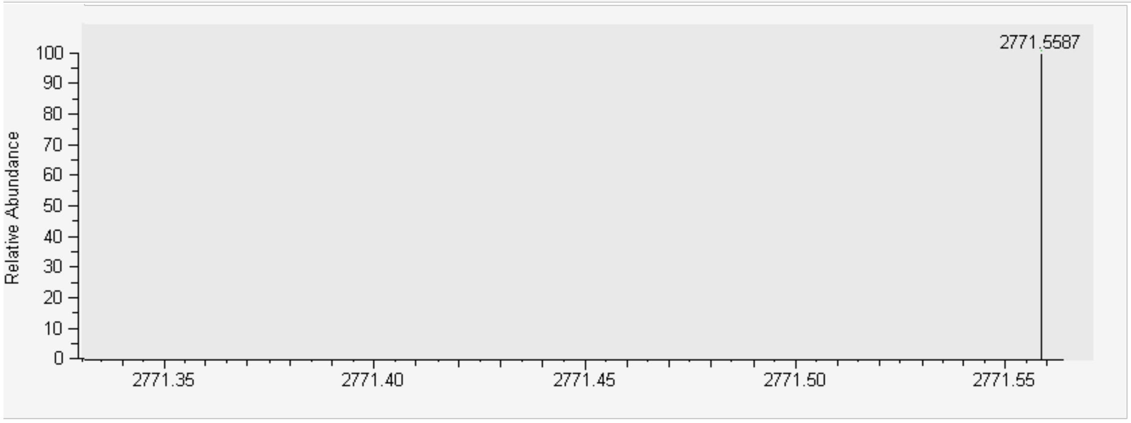

**Fig. S1. Synthesis of K15acacH2B(1-26) peptide.**

**(A)** Scheme of chemical synthesis of the 2,5-dioxopyrrolidin-1-yl 2-(2-methyl-1,3-dioxolan-2-yl)acetate. **(B)** Mass spectra (top) and $^1$H NMR spectrum (bottom) of ethyl 2-(2-methyl-1,3-dioxolan-2-yl)acetate. ESI-HRMS calc for $C_8H_{14}NaO_4$ $[M+Na]^+$: 197.0784 found 197.0779 $^1$H NMR (500 MHz, DMSO-*d6*) δ 3.95 (q, J = 7.1 Hz, 2H), 3.77 (s, 4H), 1.29 (s, 3H), 1.08 (t, J = 7.1 Hz, 3H). **(C)** Mass spectra (top) and $^1$H NMR spectrum (bottom) of 2-(2-methyl-1,3-dioxolan-2-yl)acetic acid. ESI-HRMS calc for $C_6H_{10}NaO_4$ $[M+Na]^+$: 169.0471 found 169.0466 $^1$H NMR (500 MHz, DMSO-$d_6$) δ 3.84 (s, 4H), 3.15 (s, 2H), 1.37 (s, 3H). **(D)** Mass spectra (top) and $^1$H NMR spectrum (bottom) of 2,5-dioxopyrrolidin-1-yl 2-(2-methyl-1,3-dioxolan-2-yl)acetate. ESI-HRMS calc for $C_{10}H_{13}NNaO_6$ $[M+Na]^+$: 266.0635 found 266.0628 $^1$H NMR (500 MHz, DMSO-$d_6$) δ 3.86 – 3.79 (m, 4H), 2.90 (s, 2H), 2.71 (s, 4H), 1.34 (s, 3H). **(E)** Structure (top) and ESI mass spectra (bottom) of the synthetic K15acac-H2B(1-26) peptide with the deconvoluted spectra below it. ESI-HRMS calc for $C_{122}H_{210}N_{36}O_{37}$ [M]+: 2771.5657 found 2771.5587.

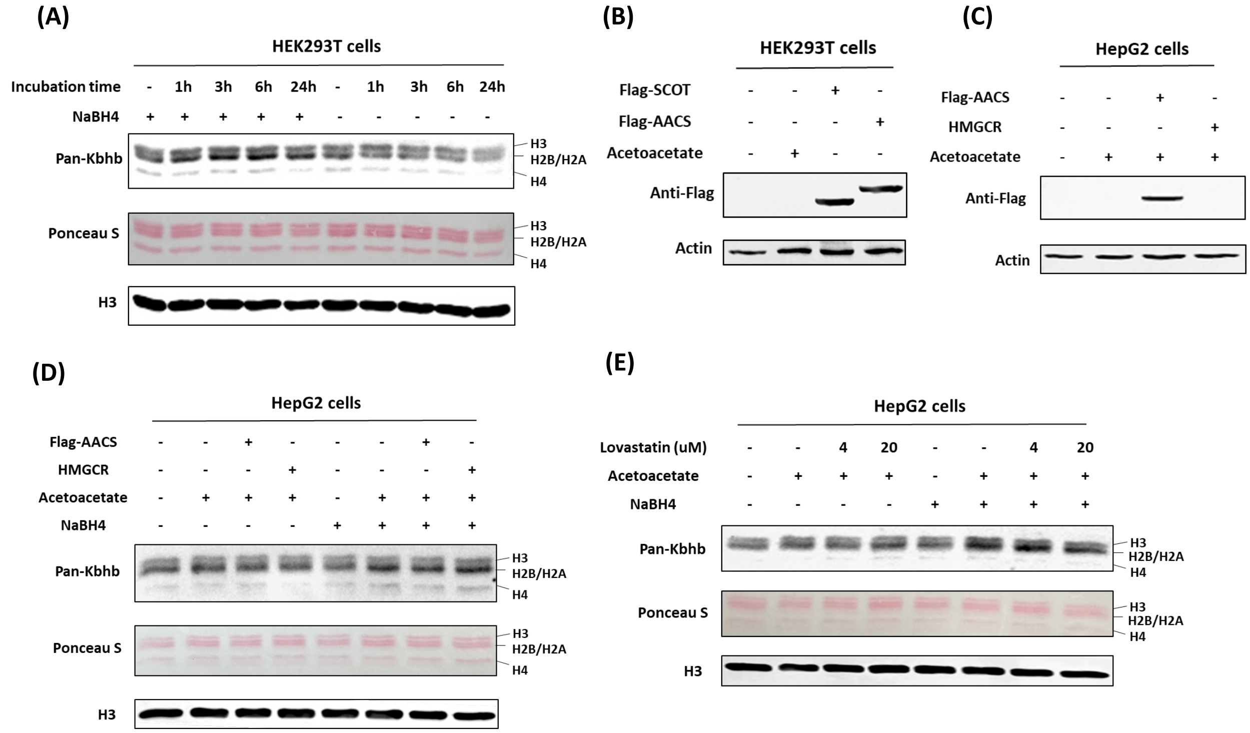


**Fig. S2. Dynamic regulation of histone Kacac in vivo.**

**(A)** Time-dependent histone Kacac in HEK293T cells treated with acetoacetate for varying durations. **(B)** Validation of SCOT and AACS overexpression in HEK293T cells. **(C)** Validation of AACS overexpression in HepG2 cells. **(D)** Western blot analysis of histone Kacac in response to AACS and HMGCR overexpression in HepG2 cells. **(E)** Western blot analysis of histone Kacac in response to treatment with lovastatin, a known HMGCR inhibitor, in HepG2 cells.

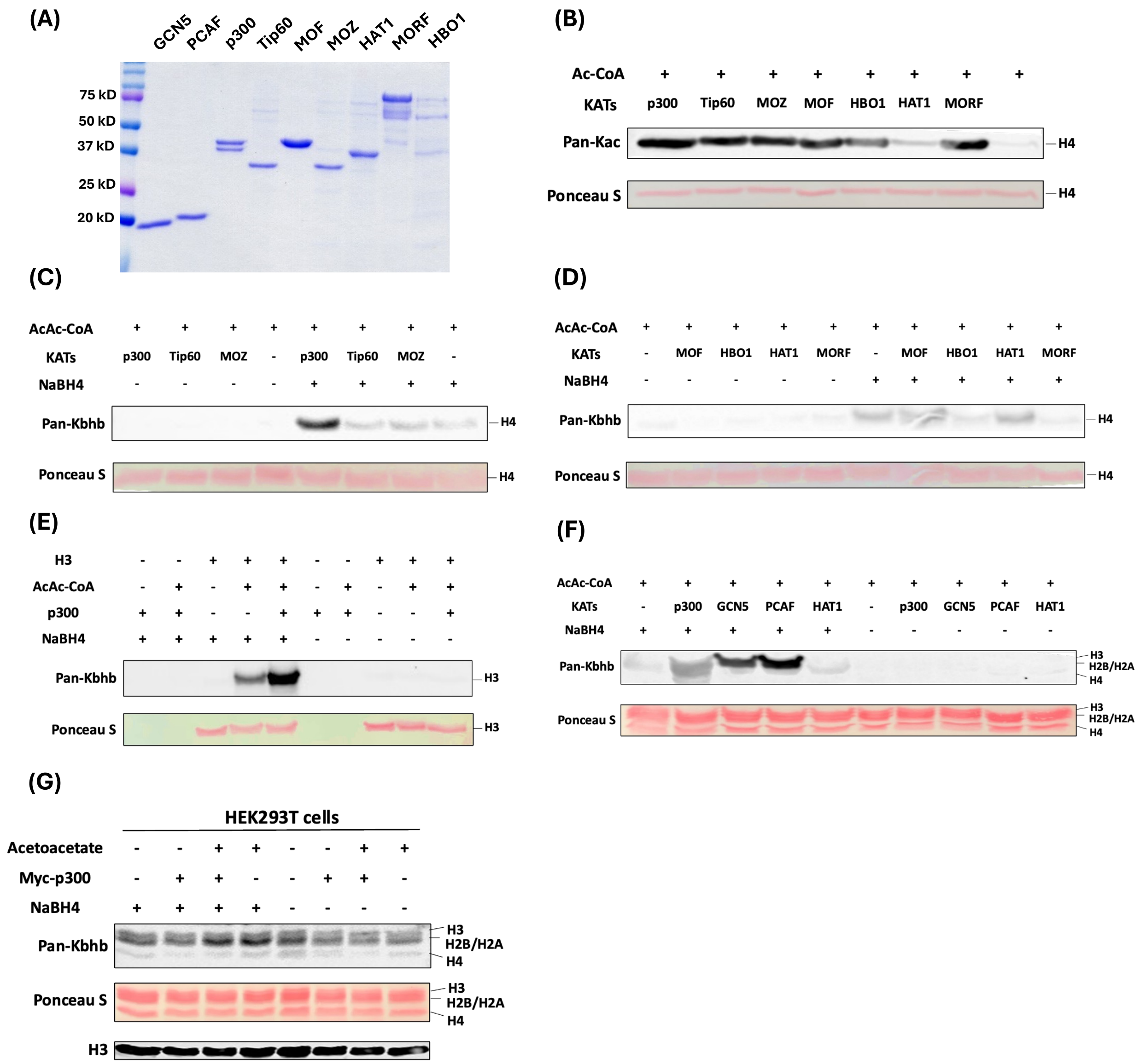


**Fig. S3. Identification of writers responsible for regulating Kacac in vitro.**

**(A)** Testing the HAT identity and purity by SDS-PAGE. **(B)** Testing the HAT activities for transferring the acetyl motif to recombinant histone H4 proteins. **(C, D)** Screening of HAT activities involved in transferring the acetoacetyl motif to recombinant histone H4 proteins. **(E)** Validation of p300-mediated Kacac on recombinant histone H3 proteins. **(F)** Validation of p300, GCN5 and PCAF activities for transferring acetoacetyl motif to histone extracts from HEK293T cells. **(G)** Western blot analysis of histone Kacac in response to p300 overexpression in HEK293T cells.

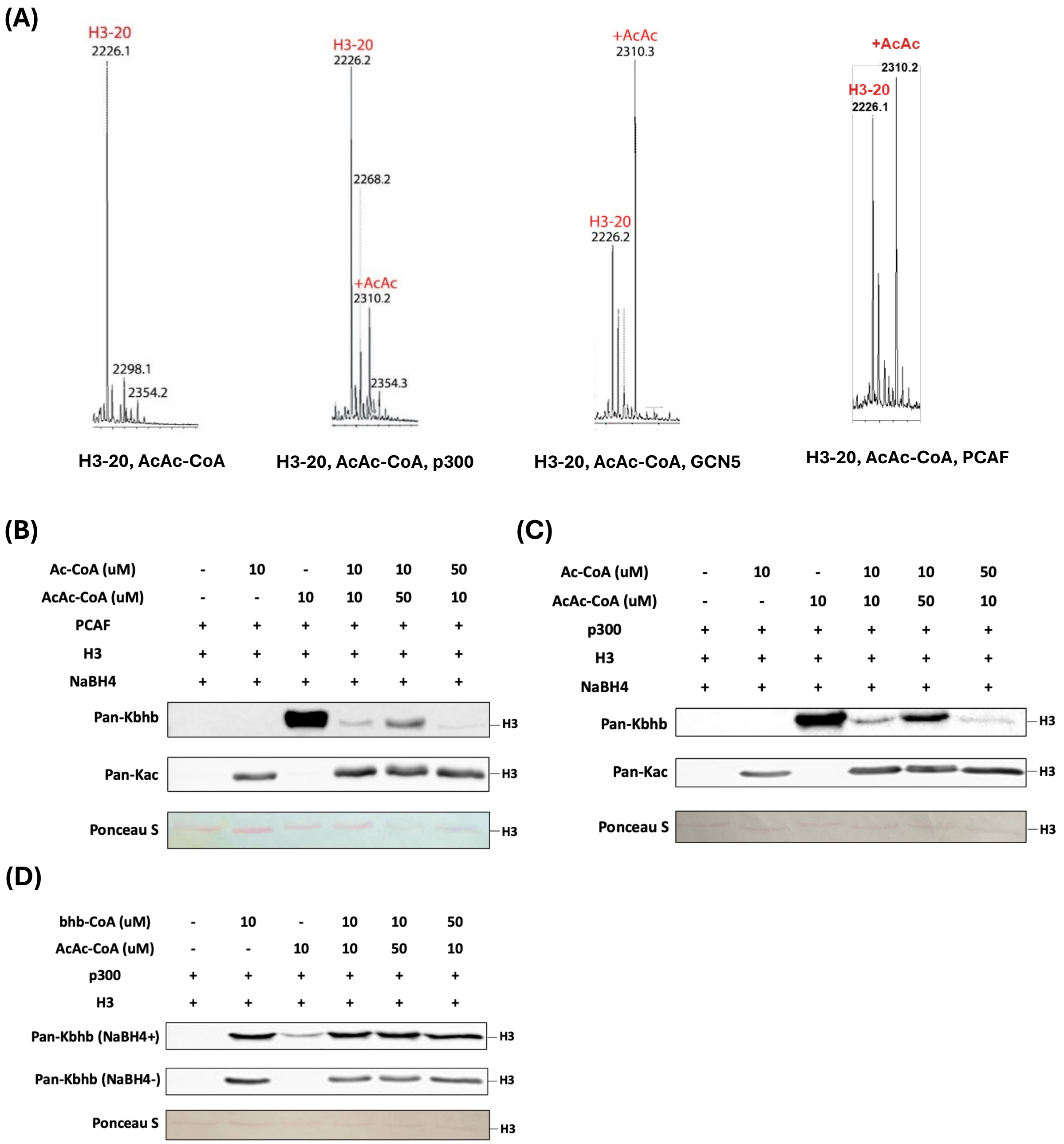


**Fig. S4. p300, GCN5 and PCAF act as acetoacetyltransferases in vitro.**

**(A)** Validation of p300, GCN5 and PCAF activities as acetoacetyltransferase on synthetic histone H3(1-20) peptide. **(B, C, D)** Proportional changes of acyl-CoAs result in dynamics of PCAF mediated substrates (B) and p300 mediated substrates (C, D) on recombinant histone H3 proteins.

**(A)**

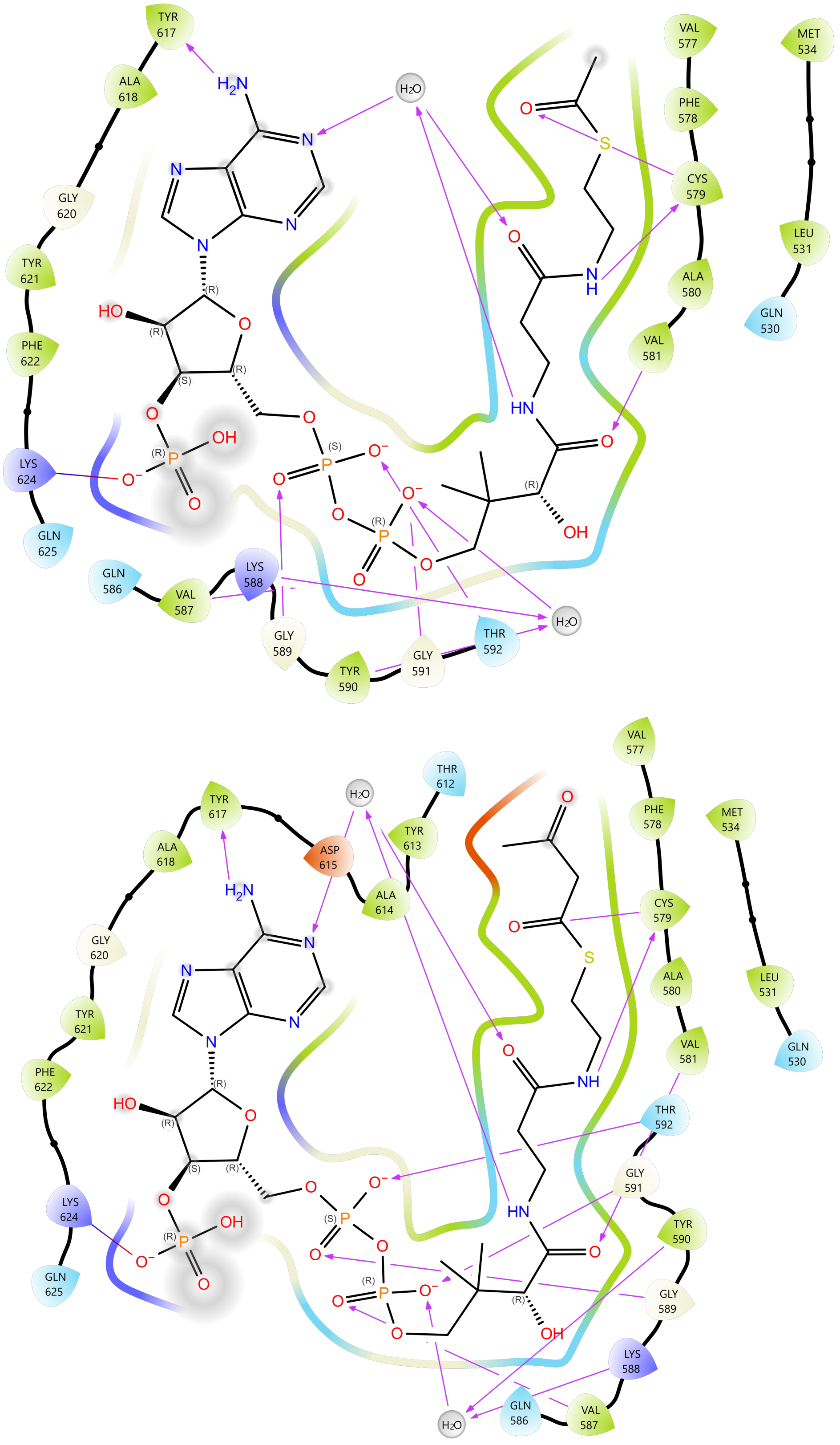
TYR 617
ALA 618
GLY 620
TYR 621
PHE 622
LYS 624
GLN 625
GLN 586
VAL 587
LYS 588
GLY 589
TYR 590
GLY 591
THR 592
VAL 577
PHE 578
CYS 579
ALA 580
VAL 581
MET 534
LEU 531
GLN 530
H2O
THR 612
TYR 613
ALA 614
ASP 615

**(B)**

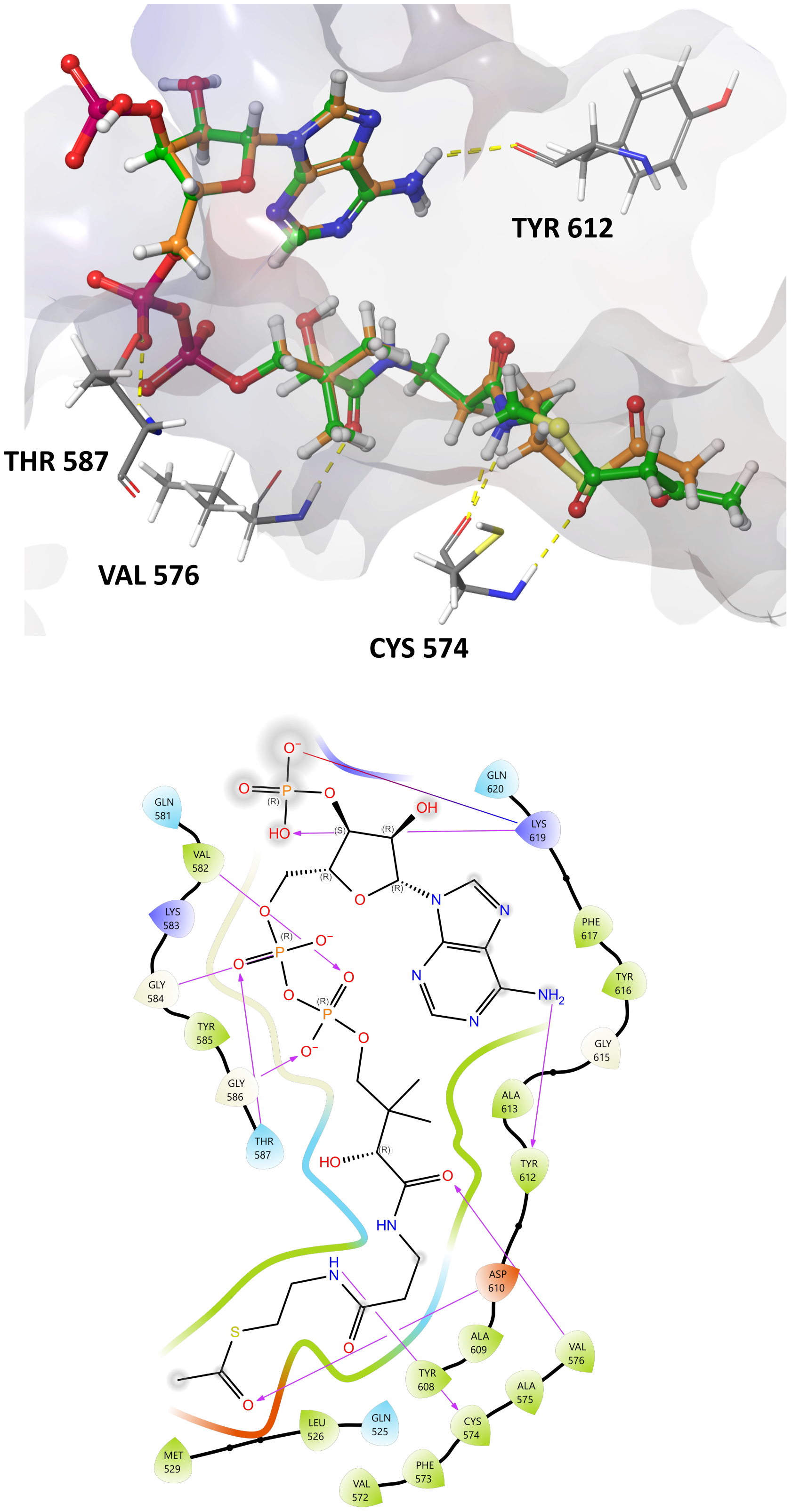
TYR 612
THR 587
VAL 576
CYS 574
GLN 620
GLN 581
LYS 619
VAL 582
LYS 583
PHE 617
TYR 616
GLY 584
TYR 585
GLY 615
GLY 586
ALA 613
THR 587
TYR 612
ASP 610
ALA 609
VAL 576
TYR 608
ALA 575
CYS 574
LEU 526
GLN 525
MET 529
PHE 573
VAL 572

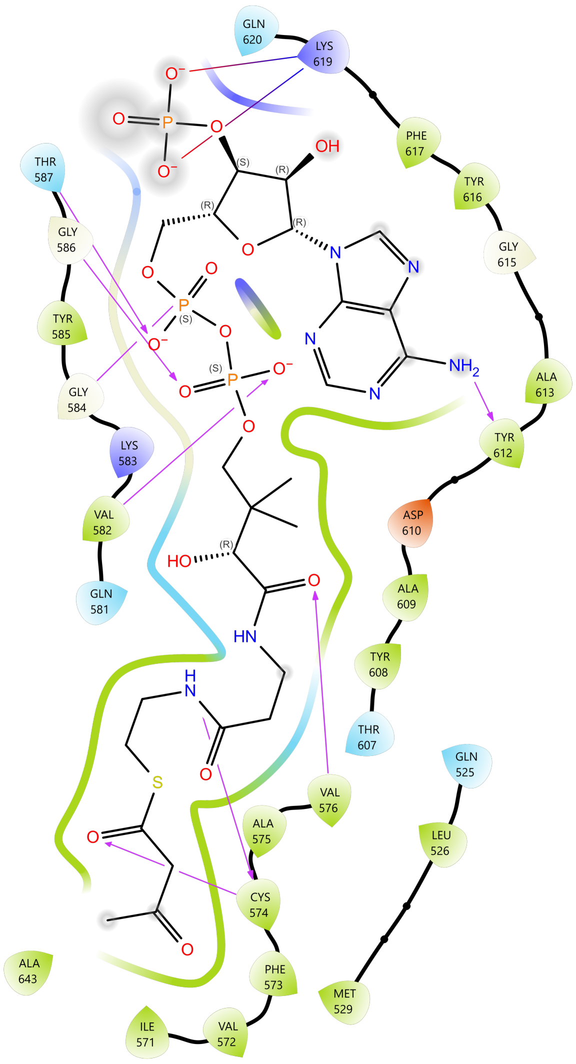
GLN 620
LYS 619
PHE 617
TYR 616
GLY 615
ALA 613
TYR 612
ASP 610
ALA 609
TYR 608
THR 607
THR 587
GLY 586
TYR 585
GLY 584
LYS 583
VAL 582
GLN 581
VAL 576
ALA 575
CYS 574
PHE 573
VAL 572
ILE 571
ALA 643
GLN 525
LEU 526
MET 529
OH
NH2
HO
HN
S

(C)

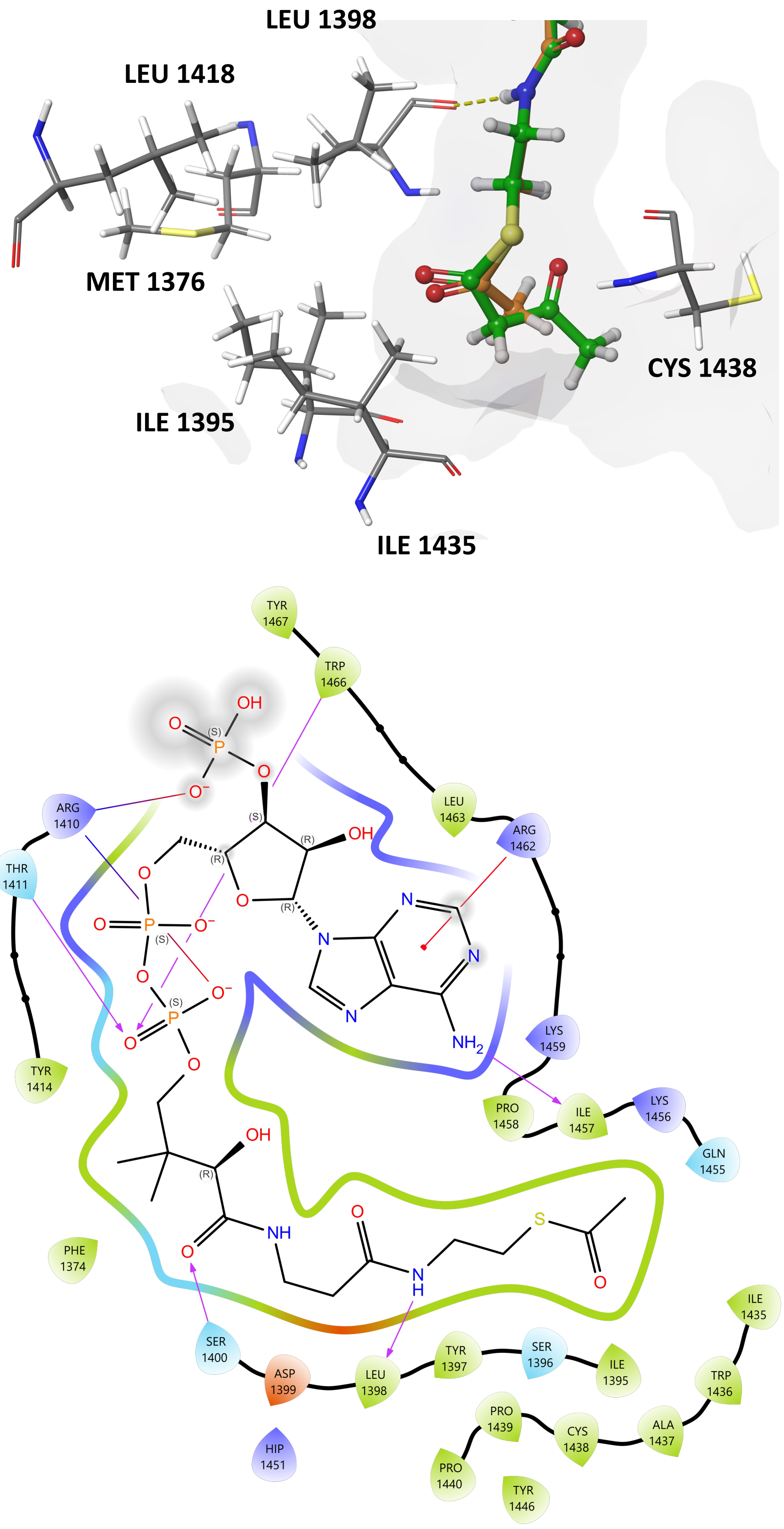
LEU 1398
LEU 1418
MET 1376
CYS 1438
ILE 1395
ILE 1435
TYR 1467
TRP 1466
LEU 1463
ARG 1462
ARG 1410
THR 1411
TYR 1414
LYS 1459
PRO 1458
ILE 1457
LYS 1456
GLN 1455
PHE 1374
SER 1400
ASP 1399
LEU 1398
TYR 1397
SER 1396
ILE 1395
TRP 1436
ILE 1435
ALA 1437
CYS 1438
PRO 1439
PRO 1440
TYR 1446
HIP 1451

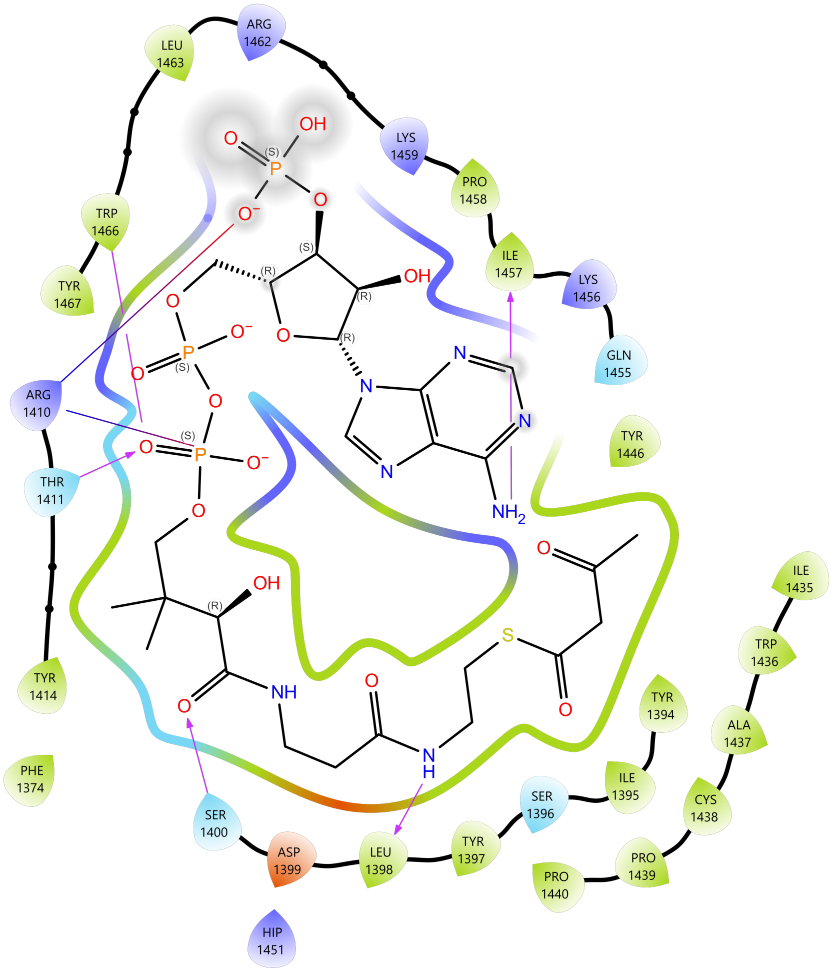
LEU 1463
ARG 1462
LYS 1459
PRO 1458
ILE 1457
LYS 1456
GLN 1455
TRP 1466
TYR 1467
ARG 1410
THR 1411
TYR 1414
PHE 1374
TYR 1446
ILE 1435
TRP 1436
TYR 1394
ALA 1437
ILE 1395
SER 1396
CYS 1438
PRO 1439
PRO 1440
SER 1400
ASP 1399
LEU 1398
TYR 1397
HIP 1451

**(D)**

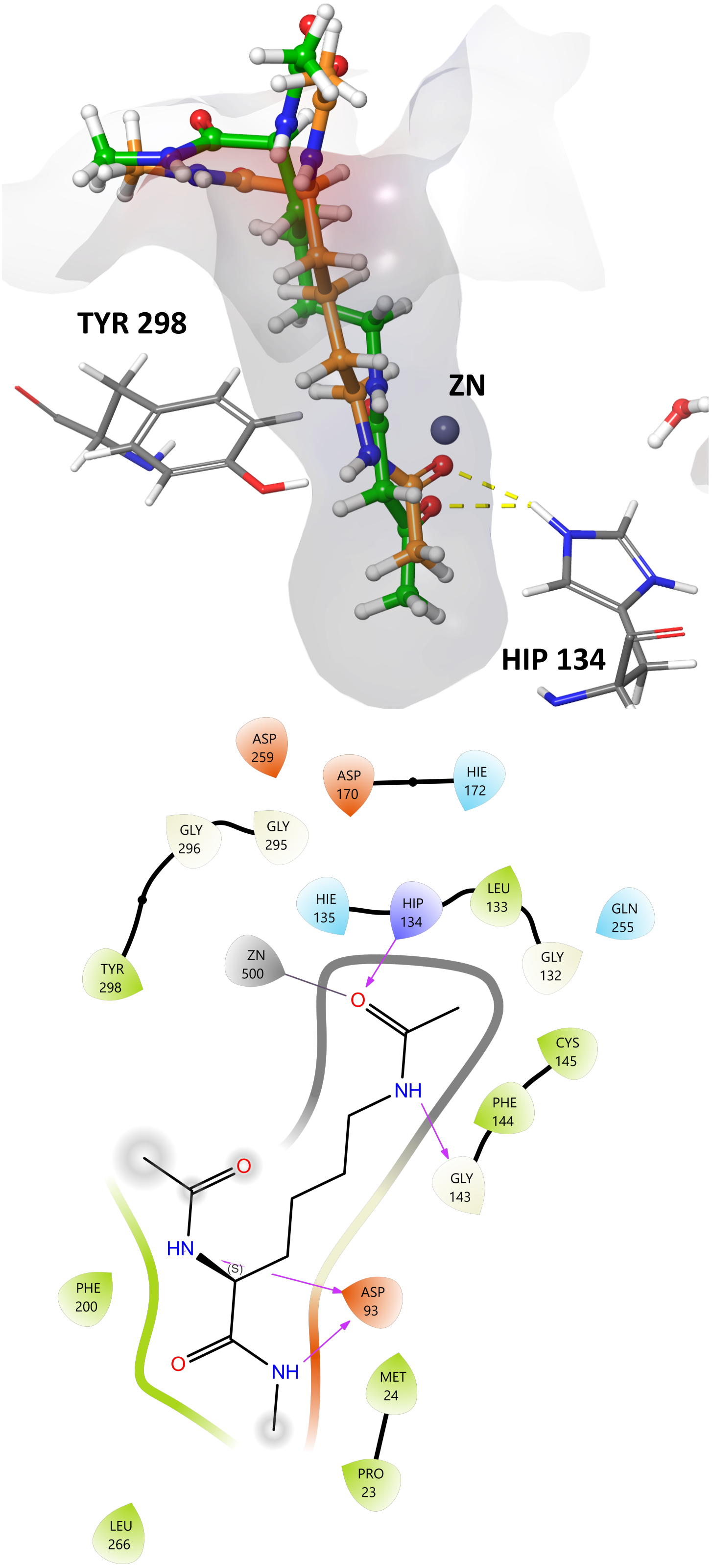
TYR 298
ZN
HIP 134
ASP 259
ASP 170
HIE 172
GLY 296
GLY 295
HIE 135
HIP 134
LEU 133
GLN 255
ZN 500
GLY 132
TYR 298
CYS 145
PHE 144
GLY 143
PHE 200
ASP 93
MET 24
PRO 23
LEU 266
O
NH
HN
(S)
O
NH

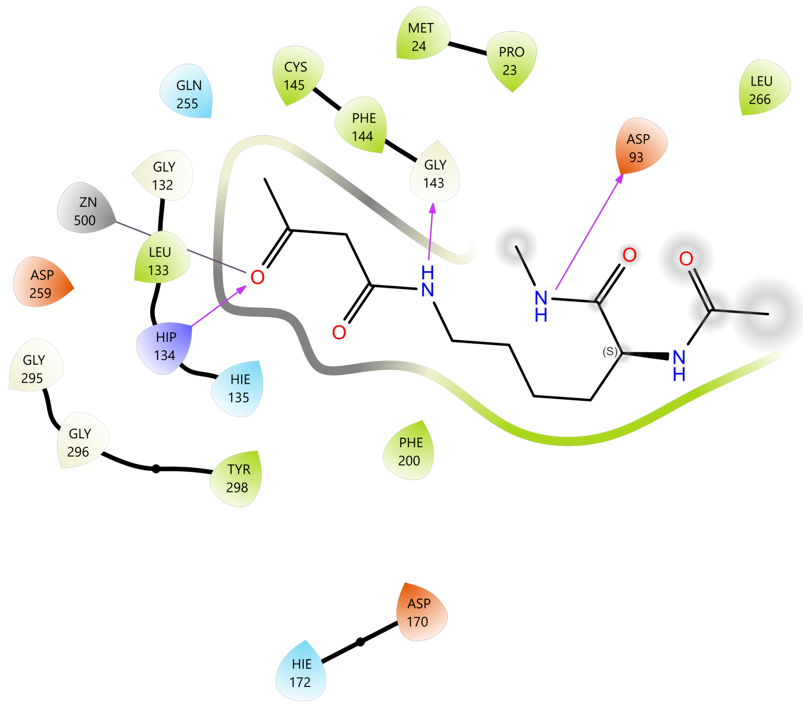


**Fig. S5. Predicted binding modes of HATs with acyl-CoAs and HDAC3 with its substrate mimics.**

**(A)** 2D diagram indicating the interaction details of GCN5 bound with acetyl-CoA (top) and acetoacetyl-CoA (bottom). PDB: 5TRL was used for the modeling. **(B)** Top: 3D diagram illustrating the catalytic pocket of PCAF bound with acetyl-CoA (orange) and acetoacetyl-CoA (green). Middle and bottom: 2D diagram indicating the interaction details of PCAF bound with acetyl-CoA (middle) and acetoacetyl-CoA (bottom). PDB: 4NSQ was used for the modeling. **(C)** Top: 3D diagram illustrating the catalytic pocket of p300 bound with acetyl-CoA (orange) and acetoacetyl-CoA (green). Middle and bottom: 2D diagram indicating the interaction details of p300 bound with acetyl-CoA (middle) and acetoacetyl-CoA (bottom). PDB: 5LKU was used for the modeling. **(D)** Top: 3D diagram illustrating the catalytic pocket of HDAC3 bound with acetyl-lysine mimic (orange) and acetoacetyl-lysine mimic (green). Middle and bottom: 2D diagram indicating the interaction details of HDAC3 bound with acetyl-lysine mimic (middle) and acetoacetyl-lysine mimic (bottom). PDB: 4A69 was used for the modeling.

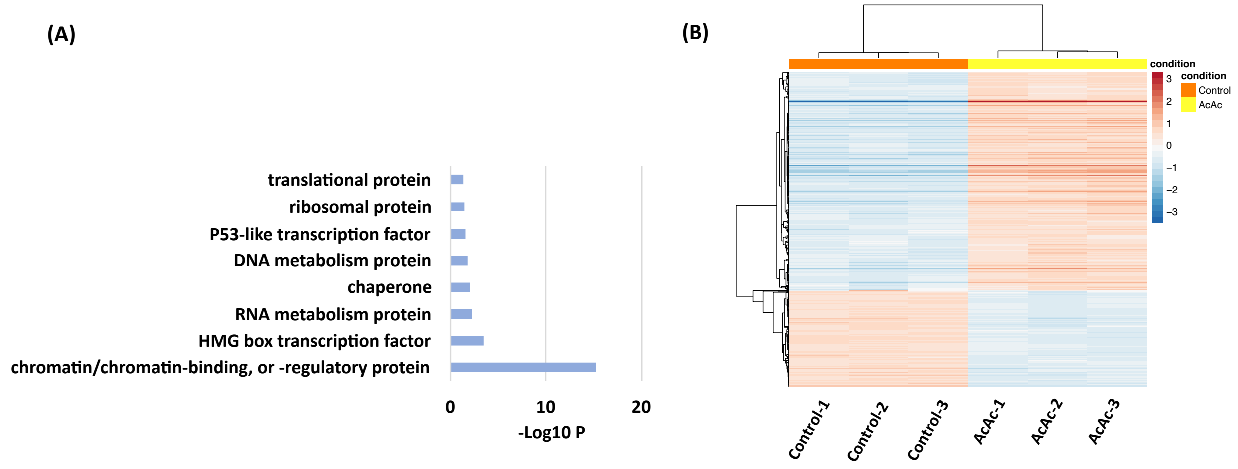


**Fig. S6. Functional annotation of Kacac marks in HEK293T cells.**

**(A)** Protein class enrichment of Kacac proteins, ranked on the basis of raw p values. **(B)** Heat map of differentially expressed genes in "Control" and "AcAc" groups in RNA-seq data. "Control" indicates untreated HEK293T cells; "AcAc" indicates lithium acetoacetate treated HEK293T cells.

**Tables S1-S6. (separate file)**

**Table S1.** Complete list of identified Kacac (DKbhb) sites.

**Table S2.** Kacac sites and neighbor sites that are critical for biological functions.

**Table S3.** Protein complex analysis of Kacac proteins.

**Table S4.** Differentially expressed genes (DEGs) upon acetoacetate treatment in RNA-seq.

**Table S5.** GO terms identified by using DEGs.

**Table S6.** KEGG pathways identified by using DEGs.
